# A Monomer-Dimer Equilibrium Tunes Phospholipid Handling by *Campylobacter jejuni* MlaC to the Bacterium’s Unique Lipidome

**DOI:** 10.64898/2026.08.28.747810

**Authors:** Liliana Fernandes da Costa, Tobias Rath, Sophie Spiewag, Lina Leipold, Christian Bonifer, Ngoc Minh Bui, Mariya Lazarova, Wuen Ee Foong, Heng-Keat Tam, Andrea Herrmann, Clemens Glaubitz, Klaas M. Pos, Nina Morgner

**Affiliations:** Institute of Biochemistry, Goethe-University Frankfurt, Max-von-Laue-Str. 9, D-60438 Frankfurt am Main, Germany; Cluster of Excellence SubCellular Architecture of Life (SCALE), Goethe University Frankfurt, Frankfurt am Main, Germany; Institute of Physical and Theoretical Chemistry, Johann Wolfgang Goethe University, Frankfurt, Germany; Institute for Biophysical Chemistry and Center for Biomolecular Magnetic Resonance (BMRZ), Goethe University Frankfurt, Max von Laue Straße 9, 60438 Frankfurt am Main, Germany

**Author notes:** Joint corresponding authors: Klaas M. Pos, Nina Morgner. Joint first authors: Liliana Fernandes da Costa, Tobias Rath. These authors contributed equally. Biochemistry Center, Heidelberg University, Im Neuenheimer Feld 328, Heidelberg, Germany (Mariya Lazarova). Department of Biochemistry and Molecular Biology, Hengyang Medical School, University of South China, Hengyang, Hunan 421001, China (Wuen Ee Foong, Heng-Keat Tam).

## Abstract

The Gram-negative bacterial cell envelope features an asymmetric outer membrane, that confers intrinsic resistance to toxins. Maintenance of this barrier relies on the Mla system, which mediates retrograde transport of mislocalized phospholipids. In *Escherichia coli*, this system comprises the lipoprotein MlaA, the periplasmic shuttle protein MlaC, and the ABC transporter complex MlaFEDB. Intriguingly, in *Campylobacter jejuni*, *mlaA* and *mlaC* share an operon with an encoded Resistance-Nodulation-cell Division antiporter potentially involved in anterograde phospholipid transport.

Here, we describe the functional and mechanistical characterization of *Cj* MlaC. Complementation experiments in *E. coli* show that *Cj* MlaC functions independently of the native Mla system. Native mass spectrometry revealed that *Cj* MlaC uniquely exists as both monomer and dimer. Lipid binding stabilized the dimer and ion mobility mass spectrometry showed that conformational transitions precede phospholipid release, suggesting a cycle between a low-affinity monomer and a higher-lipid-affinity dimer. *Cj* MlaC binds phospholipid species distinct from *Ec* MlaC, showing an increased propensity for lysophospholipids, consistent with the unusually lysophospholipid-rich lipidome of *C. jejuni,* indicative of evolutionary adaptation to this unique lipid environment. Collectively, these findings uncover structural and mechanistic features of *Cj* MlaC and support divergent physiological roles for *Cj* and *Ec* MlaC in phospholipid trafficking.

## Introduction

Antimicrobial resistance is a major global health threat, projected to cause up to 10 million deaths annually by 2050, with Gram-negative bacteria accounting for most fatal infections between 1990 and 2021^1^. This is partly due to their protective cell envelope that is composed of an inner membrane (IM) and an outer membrane (OM), encapsulating the periplasm containing a thin peptidoglycan layer. The Gram-negative bacterial OM is characterized by phospholipids (PLs) in the inner leaflet and lipopolysaccharide (LPS) molecules in the outer leaflet^2^, resulting in a defined asymmetry. The amphipathic nature of LPS results in a formidable barrier against hydrophobic compounds such as antibiotics, slowing down their diffusion across the OM^3,4^. Together with the action of envelope-spanning multidrug efflux pumps, this contributes to intrinsic multidrug resistance^5^.

Disturbances in the asymmetry of the OM lead to formation of hydrophobic patches that facilitate the uptake of hydrophobic drugs, weakening this intrinsic resistance mechanism^6^ed. To maintain the integrity and asymmetry of the OM, Gram-negative bacteria have evolved various strategies, one of which is the maintenance of lipid asymmetry (Mla) pathway^6^. The Mla pathway was first identified in *Escherichia coli* (*E*. *coli*, *Ec*) and comprises the MlaA-OmpC/F complex in the OM, the periplasmic lipid-binding protein MlaC, and the ATP-binding cassette (ABC) transporter complex MlaFEDB in the IM, which assembles with a stoichiometry of 2:2:6:2^7,8^. The Mla pathway was thereby proposed to mediate retrograde phospholipid transport by retrieving mislocalized phospholipids from the outer leaflet of the OM and returning them to the IM^6,9^. Regarding its mechanism, molecular dynamics simulations suggest that the outer membrane component MlaA sequesters mislocalized phospholipids by interacting with their polar head groups^10,11^. The PLs are then delivered through the amphipathic channel of MlaA to the periplasmic shuttle protein MlaC, which is recruited via electrostatic interactions with MlaA^12,13^. The periplasmic shuttle protein MlaC possesses a hydrophobic pocket that accommodates the acyl chains of the phospholipids while leaving the polar head group exposed to the aqueous environment^8,14^. Following affinity-driven binding of phospholipids, MlaC transfers the phospholipids to the MlaFEDB complex, where they are incorporated into the inner membrane under ATP consumption^8,15–17^.

Among different Gram-negative bacteria, the genomic organization of the Mla system varies. While in some species, such as *Neisseria gonorrhoeae*, *Neisseria meningitidis*, and *Chlamydia* species, the Mla genes are clustered within a single operon^18,19^, in *E*. *coli*, the *mlaA* gene is genomically separated from the *mlaFEDCB* operon^6^. Curiously, *Campylobacter jejuni* (*C*. *jejuni, Cj*), a major cause of bacterial gastrointestinal infections^20^, possesses a lipidome in which lysophospholipids (lyso-PLs) account for 28-45% of all phospholipids^21^. Notably, lyso-PLs have been linked to bacterial motility and the latter to colonization propensity of the cecum in chicken^22^. On the *C. jejuni* genome, *mlaA* and *mlaC* share an operon with an encoded Resistance-Nodulation-cell Division (RND) transporter, which shares high sequence identity with the lipid transporter MmpL3 from *Mycobacterium tuberculosis* (Figure S1A) ^23^. Given the operonic co-localization and that RND transporters function usually as proton (H^+^) or sodium (Na^+^)/drug antiporters^24,25^, we hypothesize that the *C*. *jejuni* Mla pathway might function in anterograde direction, although retrograde transport cannot be ruled out at this point (Figure S1B). In order to elucidate the mechanisms of phospholipid binding, we functionally and mechanistically characterized *Cj* MlaC, revealing its phospholipid-binding properties and the mechanism underlying phospholipid transfer. Our findings reveal that *Cj* MlaC can form a dimeric, structurally distinct variant of MlaC and suggest a role in a periplasmic phospholipid and lyso-PL transfer mechanism different from that mediated by the monomeric *Ec* MlaC.

## Material and Methods

### Bacterial strains

Functional characterization of *Cj* MlaC expressed from the pTTQ18 vector was carried out in *E*. *coli* BW25113 strains. The *E. coli* BW25113 *mlaA::kan*, *mlaC::kan*, *mlaD::kan*, and *mlaE::kan* knockout strains were obtained from the KEIO collection^26^. The kanamycin resistance cassettes were removed by FLP recombination expressed from the pCP20 plasmid as described previously^27^. *Cj* MlaC and *E*c MlaC were overproduced in *E*. *coli* BL21(DE3) and the sybody Sy7J6 in *E*. *coli* MC1061.

### Cloning

The genes encoding the full-length *Cj* MlaC (*Cj*1372) and *Ec* MlaC with a C-terminal His_7_ tag were PCR-amplified from the genomic DNA of *C. jejuni* subsp. *jejuni* NCTC 11168 = ATCC 700819 and *E*. *coli* K-12 BW25113, respectively, and cloned into the pTTQ18 expression vector^28^ via Gibson Assembly^29^. The signal peptide-cleaved form of *Cj* MlaC (*Cj*1372, residues 17–189) encoded on the pTTQ18 vector was generated by inverse PCR using primers flanking the mature coding region. The PCR product was subsequently phosphorylated and ligated using the T4 DNA ligase (Thermo Fisher Scientific) prior to transformation. For in-gel GFP-based expression analysis, the His_7_ tag on the pTTQ18 vector encoding full-length *Cj* MlaC was replaced with a C-terminal superfolder green fluorescent protein (sfGFP) tag by Gibson assembly, using the p7XC3GH vector as template^30^. For overexpression, the gene encoding the signal peptide-cleaved form of *Cj* MlaC with an N-terminal His_6_ tag and a C-terminal Avi-tag was PCR-amplified from the genomic DNA and cloned into the pET24a vector (Novagen) via Gibson Assembly. In turn, the signal peptide-cleaved form of *Ec mlaC* (residues 22–211) was PCR-amplified from the genomic DNA using primers containing NdeI and XhoI restriction sites and cloned into the pET24a vector resulting in the coding of *Ec* MlaC with a C-terminal His_6_ tag. The resulting plasmids were propagated in *E. coli* DH5α cells and were verified by restriction enzyme digestion and Sanger sequencing. Primer sequences are listed in Table S1. The pSBinit plasmid encoding the sybody Sy7J16 with a C-terminal c-Myc tag and His_6_ tag was obtained from a synthetic sybody library^31^.

### Growth curve analysis

A single colony of cells transformed with the respective pTTQ18 vector was grown overnight in lysogeny broth (LB) medium supplemented with 50 µg/mL carbenicillin at 37 °C with shaking at 130 rpm. On the following day, the overnight culture was diluted into fresh LB medium without antibiotics to an initial optical density (OD_600_) of 0.018. Then, 100 µL of LB medium containing 50 µg/mL carbenicillin and the antimicrobial agent to be tested was transferred into each well of a sterile 96-well plate and mixed with 50 µL of the diluted culture. The plate was incubated at 37 °C for 20 hours in an EON microplate reader (BioTek) and OD_600_ measurements were automatically performed every 20 min. All experiments were conducted in at least three independent biological replicates. The growth curves were illustrated using Origin 2023b (OriginLab Corporation).

### Western blot analysis of expressed MlaC

To assess the protein levels of MlaC expressed from the pTTQ18 vector in *E. coli* BW25113, Western blotting was performed. For this, LB medium supplemented with 50 µg/mL carbenicillin was inoculated with a single colony carrying the respective pTTQ18 construct. Cultures were grown at 37 °C and 130 rpm for 20 h. Cells were harvested and resuspended at an OD_600_ of 20 in 1 mL, followed by lysis in 100 µL of lysis buffer (20 mM Tris pH 8.0, 150 mM NaCl, 10 µg/mL lysozyme, 10 µg/mL DNase I, and 200 µM PMSF). Lysis was performed using a FastPrep machine at 2,650 rpm and 4 °C for 4 °C with glass beads. Lysates were afterwards clarified by centrifugation at 13,000 x *g* for 10 min at 4 °C, and 15 µL of the supernatant were subsequently mixed with 5X SDS loading dye (250 mM Tris-HCl pH 6.8, 5% (v/v) β-mercaptoethanol, 0.02% (w/v) bromophenol blue, 30% (v/v) glycerol, 10% (w/v) SDS) and analyzed by sodium dodecyl sulfate–polyacrylamide gel electrophoresis (SDS-PAGE) and subsequent Western blotting. Membranes were blocked with 3% BSA in TBS-T (20 mM Tris-HCl, 150 mM NaCl, 0.1% Tween-20, pH 7.6) for 1 h at room temperature. Afterwards, the blots were incubated overnight at 4°C with mouse anti-His_6_ primary antibody (Abcam, ab1187) diluted 1:1,000 in 3% BSA in TBS-T, followed by three washes with TBS-T for 5 min each. Subsequently, membranes were incubated for 1 h at room temperature with goat anti-mouse alkaline phosphatase (AP)-conjugated secondary antibody (Sigma Aldrich, A5153) diluted 1:2,000 in 1% BSA in TBS-T, followed by three washes with TBS-T. The protein signals were developed using the 5-bromo-4-chloro-3-indolyl phosphate (BCIP) and nitro-blue tetrazolium (NBT) substrates diluted in AP buffer (100 mM Tris pH 9.5, 10 mM MgCl_2_, 100 mM NaCl). Expression of full-length *Cj* MlaC with a C-terminal sfGFP tag was additionally assessed by in-gel sfGFP fluorescence. Samples were prepared as described for the Western blot analysis, separated by SDS-PAGE, and imaged using a Fusion Fx imaging system, Vilber (λ_ex_/ λ_em_ =488 nm/509 nm).

### Expression and Purification

For large-scale expression, a pre-culture of LB medium (10 g/L tryptone, 5 g/L yeast, 10 g/L NaCl) containing 50 µg/mL kanamycin was inoculated with a single colony of *E*. *coli* BL21(DE3) transformed with the pET24a vector encoding either *Cj* MlaC or *Ec* MlaC. For overexpression of the sybody Sy7J6, *E*. *coli* MC1061 cells were transformed with the plasmid pSBinit_Sy7J6_c-Myc_His_6_-tag, and a pre-culture was prepared in LB medium supplemented with 25 µg/mL chloramphenicol. After overnight incubation at 37 °C with shaking at 130 rpm, 1 L of Terrific Broth (TB) medium (12 g/L tryptone, 24 g/L yeast extract, 4 mL/L (v/v) glycerol, 2.31 g/L KH₂PO₄, 12.54 g/L K₂HPO₄), supplemented with the corresponding antibiotic, was inoculated with 10 mL of the overnight culture and incubated at 37 °C with shaking at 130 rpm until an optical density at 600 nm (OD_600_) of 0.5–0.6 was reached. Gene expression was then induced with 1 mM isopropyl β-D-1-thiogalactopyranoside (IPTG) overnight at 20 °C for *Cj* and *E*c MlaC, or with 0.2% (v/v) arabinose for the sybody. On the following day, the cells were harvested by centrifugation at 17,600 x *g* for 20 min and resuspended in 2 mL of lysis buffer (20 mM Tris pH 8.0, 150 mM NaCl, and 20 mM imidazole, 10 µg/mL lysozyme, 10 µg/mL DNase I, and 200 µM PMSF) per gram of cells. The cell suspension was stirred for 40 min and then lysed by two passes through a Stansted SPCH-EP-10 530 pressure cell homogenizer (Homogenizing Systems Ltd.) at 22 kPsi. Cell debris and membranes were removed by centrifugation at 32,000 x *g* for 1 h at 4 °C. The supernatant, containing the His-tagged soluble proteins, was then passed twice over nickel– nitrilotriacetic acid (Ni-NTA) resin pre-equilibrated with lysis buffer. Subsequently, the resin was washed twice with 15 column volumes of lysis buffer, and the proteins were eluted with 10 column volumes of lysis buffer containing 250 mM imidazole and 0.1 mM ethylenediaminetetraacetic acid (EDTA). The eluted proteins were concentrated using an Amicon Ultra-15 centrifugal filter unit (10 kDa molecular weight cutoff) and buffer-exchanged by size-exclusion chromatography (SEC) using a Superose 6 10/300 Increase column equilibrated in 20 mM Tris pH 8.0, 150 mM NaCl, at a flow rate of 0.3 mL/min. All purification steps were performed at 4 °C. Protein purity was assessed by SDS-PAGE followed by Coomassie staining.

### Lipid extraction from purified protein

Co-purified lipids were extracted from MlaC as previously described^32^. Briefly, 2 mL of purified protein (1 mg/mL) was mixed with 2 mL of methanol and 1 mL of chloroform (2:2:1, v/v). The mixture was vortexed for 5 min, incubated at 50 °C for 30 min, and vortexed again for 5 min. The samples were then centrifuged at 2,000 × *g* for 10 min, and the lower organic phase was transferred to a fresh glass tube using a glass Pasteur pipette. Solvent was evaporated at 100 °C in a water bath, and the dried lipid film was resuspended in 100 µL of chloroform.

For thin-layer chromatography (TLC) analysis, 20 µL of the extracted lipid samples and 5 µL of *E*. *coli* total lipid extract (Avanti Polar Lipids, 100600) as positive control were applied to a silica TLC plate (Carl Roth, N726.1). As negative control, lipid extracts from a sybody were used. The extracted lipids were separated by TLC using a solvent system consisting of chloroform:methanol:acetic acid (65:25:10, v/v/v). Afterwards, the plate was air-dried for 30 min and visualized by iodine vapor staining in a sealed chamber containing iodine crystals.

### Delipidation of *Cj* MlaC

For delipidation of MlaC, cell lysates with expressed *Cj* MlaC were passed twice through a pre-equilibrated Ni-NTA affinity column and washed twice with 25 mL of detergent buffer (20 mM Tris, pH 8.0, 150 mM NaCl, 20 mM imidazole, 25 mM *n*-nonyl-β-D-glucopyranoside (β-NG) (Anatrace, N324) by static incubation for 1 hour. This was followed by an additional overnight wash under gentle rotation using the same buffer. A final wash step was performed with 25 mL of lysis buffer (20 mM Tris, pH 8.0, 150 mM NaCl, 20 mM imidazole) for 1 hour by static incubation. The protein was then eluted from the Ni-NTA column with 10 column volumes of 20 mM Tris, pH 8.0, 150 mM NaCl, 250 mM imidazole, and 0.1 mM EDTA. The protein was concentrated using an Amicon Ultra-15 centrifugal filter unit (10 kDa MWCO; Merck Millipore) and buffer-exchanged by SEC using a Superose 6 10/300 Increase column to 20 mM Tris, pH 8.0, 150 mM NaCl. All purification steps were performed at 4 °C. Protein purity was assessed by SDS-PAGE followed by Coomassie staining.

### 31P nuclear magnetic resonance (NMR) spectroscopy

^31^P NMR spectra were acquired on a 400 MHz spectrometer (Bruker) with a 5 mm BBO probe with Z-gradient (Bruker), operating at a ^31^P frequency of approximately 162 MHz. Spectra were recorded in direct detection mode with 17.8 W Waltz-65 ^1^H decoupling. For each sample, 500 µL of 20 mg/mL *Cj* MlaC protein (lipidated or delipidated) or of the first detergent wash fraction were mixed in a 5 mm NMR tube with 50 µL of a 3 mM stock solution of 2,2-dimethyl-2-silapentane-5-sulfonate (DSS) in D_2_O. Chemical shifts were referenced to DSS at 0 ppm. Data were acquired at 298 K using 8,192 experimental scans with a recycle delay of 2 s. Free induction decays (FIDs) were processed using TopSpin 4.0.9 (Bruker), applying exponential line broadening of 5 Hz.

### Nano electro spray ionization mass spectrometry (nESI-MS)

All nESI experiments were conducted using a Synapt G2-S instrument (Waters Corporation, Wilmslow, UK). MlaC samples were prepared by a buffer exchange into freshly prepared 200 mM ammonium acetate using Amicon Ultra 0.5 centrifugal filters with a cutoff of 10 kDa. As nESI source, gold-coated borosilicate capillaries were used, which were pulled by P1000 Flaming/Brown Micropipette Puller (Sutter Instrument, Novato USA) before sputtering with gold. 5 µL of 10 µM sample solution were loaded directly into the capillary, and 1.6 kV were applied to the needle. The cone voltage was set to 100 V, while the source temperature was at 30 °C. Optimized for high m/z ratios, the detector was calibrated using a conventional CSI calibration. The transfer collision voltage was varied between 0 and 150 V, while the trap collision voltage was set between 2 and 5 V. Spectra were obtained with a scan rate of 1 s^-1^. All measurements were conducted in the positive ion mode.

Lipid extracts were measured directly from chloroform, loading 10 µL into the capillary. The capillary voltage was adjusted to 2.0 kV, and between 50 and 100 V of transfer collision voltage was used. The major phospholipid peaks absent in the chloroform reference spectrum were identified using the MAPS® lipid database^33,34^.

*In vitro* lipidation studies with delipidated *Cj* MlaC were done at a 1:5 protein-to-lipid molar ratio. The phospholipids (Table S2) were resuspended in 200 mM ammonium acetate (pH 7.0) containing 0.4% (v/v) β-NG. Following incubation for 1 h at room temperature, samples were diluted to a final protein concentration of 10 µM prior to nESI-MS analysis.

For Tandem MS/MS experiments, the standard G2-S quadrupole filter was used for ion isolation. Collision induced dissociation was performed after precursor selection from 10 – 110 V of collision energy in the transfer cell.

Ion mobility spectrometry (IMS) measurements were conducted using a step wave setup with the following parameters. The wave height was 40 V, with a traveling wave velocity of 700 m/s. The drift cell pressure was at 5.7 mbar, while the nitrogen gas flow was set to 70 mL/min. The collision ramp was conducted from 0-100V of transfer collision energy (CE), using steps of 5 V.

For data analysis of MS, MS/MS and IMS data, the software MassLynx, TWIMExtract ^35^, UniDec ^36^ and OriginPro 2022b (Origin Lab Corporation) were used.

### Laser induced liquid bead ion desorption mass spectrometry (LILBID-MS)

LILBID-MS was carried out with purified samples according to the protocols described in^37^. In short, all protein samples were buffer exchanged to 200 mM ammonium acetate using Zeba spin desalting columns. These samples were then diluted to 10 µM and loaded directly into a piezo-driven droplet generator (MD-K-130, Microdrop Technologies GmbH). At a frequency of 10 Hz, droplets of a diameter of around 50 µm are transferred into a high vacuum, where they are irradiated by a pulsed IR Laser, which works at a wavelength of 2.8 µm, exciting the asymmetric O-H stretch vibration of water. This leads to an explosive expansion of the water droplets, setting the ions free, which are then accelerated, via an applied acceleration voltage, in the Wiley McLaren type ion optics, into the homebuilt time-of-flight analyzer. Through a reflectron the ions are then guided into the Daly type detector, which is optimized for analysis of high m/z ratios. The voltages of repeller and extractor of the ion optics were set to – 4.0 kV each, while the latter was pulsed to – 6.6 kV for 370 µs, 10-20 µs after droplet explosion. The reflectron was set to -7.2 kV, while the Einzel lenses in front of it were set to -3.0 kV each. Spectra processing was done by OriginPro 2023b (Origin Lab Corporation).

## Results

### *Cj* MlaC functions independently of the *E. coli* Mla pathway

*Cj* MlaC shares 21.6% sequence identity with MlaC from *E*. *coli,* the latter identified as part of the Mla system^6^. The absence of *Ec* MlaC in *E*. *coli* has been shown to enhance susceptibility against toxins, such as doxycycline, as a result of the compromised retrograde lipid transport from the outer leaflet of the OM^6^. To investigate whether *Cj* MlaC can complement the role in retrograde phospholipid transport in *E*. *coli* BW25113 Δ*mlaC*, a plasmid encoding *Cj* MlaC was tested for mitigation of the growth defect in presence of the antibiotic doxycycline. The presence of the *Cj mlaC* gene was able to restore *E*. *coli* Δ*mlaC* growth in the presence of doxycycline in a similar fashion as the introduction of the *Ec mlaC* gene (Figure 1A). Intriguingly, however, the presence of the *Cj mlaC* gene also markedly improved bacterial growth under doxycycline exposure in strains devoid of genes encoding the other components of the retrograde lipid transfer Mla system (Δ*mlaA*, Δ*mlaD* or Δ*mlaE*). Although depletion of *mlaE* had an especially strong detrimental effect on bacterial growth, *Cj* MlaC nevertheless was able to improve the bacterial growth (Figure 1). In contrast, the *Ec mlaC* gene could not rescue the reduced growth phenotype other than that of *E*. *coli* Δ*mlaC*. The rescue phenotype of *Cj* MlaC was dependent on its periplasmic localization, since the variant lacking the coding sequence for the periplasmic signal sequence could not complement the deletion of *Ec mlaC* in the doxycycline-sensitive phenotypes. To our surprise, complementation with *Cj mlaC* also improved growth of *E*. *coli* BW25113 Δ*mlaA*, Δ*mlaC* or Δ*mlaD*, but not the growth of *E*. *coli* BW25113 Δ*mlaE* on LB medium without antibiotics (Figure S2A). These findings suggest that the presence of *Cj* MlaC provides an overall fitness benefit to *E. coli*, which is pronounced under antibiotic stress. To rule out the possibility that the observed doxycycline resistance is specific for this antibiotic for example due to direct binding of the antibiotic to *Cj* MlaC in the periplasm, the growth curve analyses were repeated with vancomycin, a large and highly polar glycopeptide that cannot readily cross the *E. coli* outer membrane and is not anticipated to bind to *Cj* MlaC^38^. Again, the presence of *Cj* MlaC provided a growth-promoting effect in the presence of vancomycin, as was observed under doxycycline exposure (Figure S2B). This supports the hypothesis that the resistance was not caused by direct antibiotic binding to *Cj* MlaC in the periplasm but rather hinted toward the restoration of the OM asymmetry.

**Figure 1.**
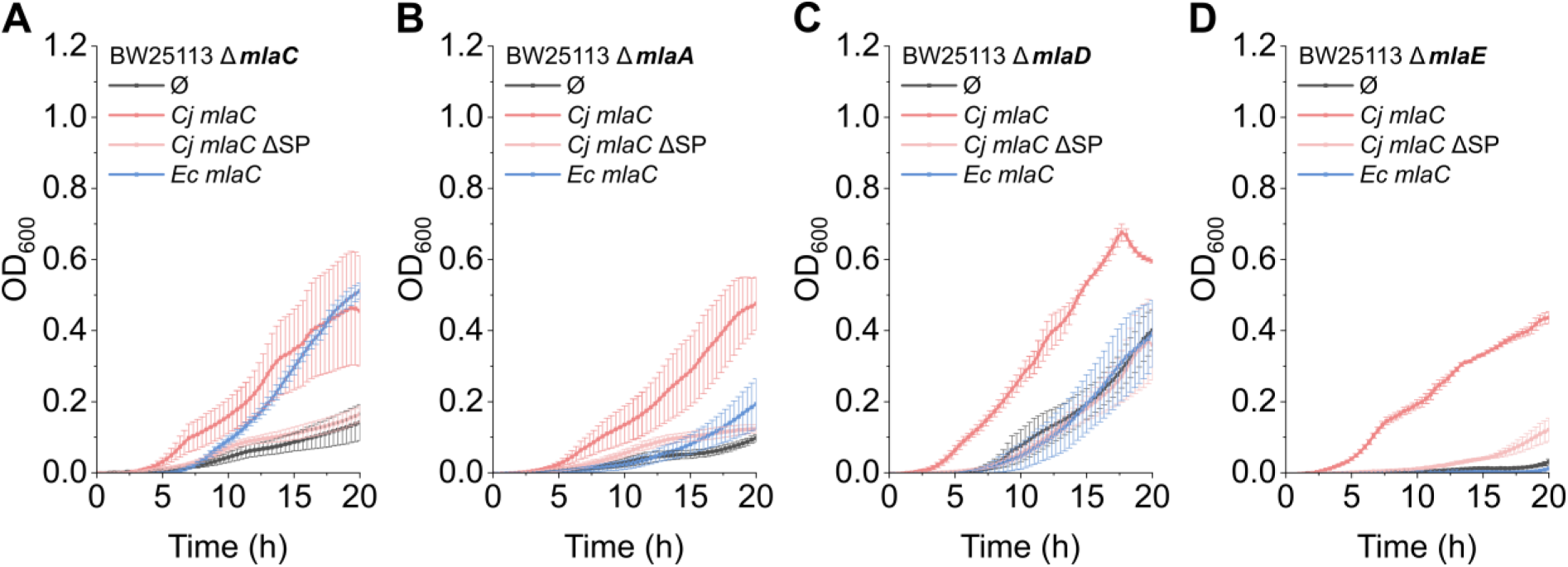
Functional characterization of *Cj* MlaC in *E*. *coli*. (**A-D**) Growth curve analysis of *E*. *coli* (*Ec*) BW25113 cells lacking the endogenous *mlaC*, *mlaA*, *mlaD*, or *mlaE* genes, complemented with either the empty pTTQ18 vector (gray, Ø), pTTQ18 encoding *C*. *jejuni* (*Cj*) MlaC (dark red), the signal-peptide cleaved *Cj* MlaC variant (light red, *Cj mlaC* ΔSP) or *Ec* MlaC (blue). Cells were grown in lysogeny broth (LB) containing 50 µg/mL carbenicillin as selection antibiotic and 0.4 µg/mL doxycycline as antimicrobial agent for 20 h at 37 °C. Optical density at 600 nm (OD600) was measured at the indicated time points and plotted over time. Each data point is presented as the mean ± standard error of the mean (SEM) from three biological replicates.

Western blot analyses indicated that *Cj* MlaC is poorly expressed in *E. coli*, in contrast to *Ec* MlaC, while the *Cj* MlaC variant lacking the periplasmic signal sequence was expressed at levels similar to *Ec* MlaC (Figure S3A). To confirm expression of full-length *Cj* MlaC, protein levels of sfGFP-tagged *Cj* MlaC in the BW25113 wild-type strain were assessed by in-gel GFP fluorescence, as fluorescence detection is more sensitive than Western blotting. sfGFP-tagged *Cj* MlaC was readily detected (Figure S3B), confirming expression of the full-length fusion protein. Furthermore, the sfGFP-tagged MlaC construct was functional, as demonstrated by complementation of the *E*. *coli* BW25113 Δ*mlaC* strain, resulting in an enhanced growth phenotype in presence of doxycycline (Figure S3C), similar to the result obtained by complementation with the full-length His-tagged *Cj* MlaC (Figure 1A). This suggests that the sfGFP-tagged *Cj* MlaC fusion protein is successfully transported to the periplasm. Despite its lower abundance compared to its *E. coli* counterpart, periplasmically localized *Cj* MlaC markedly enhanced bacterial growth, particularly under antibiotic stress, indicating that it is functionally more effective than *Ec* MlaC. Together, these findings suggest that the pronounced activity of *Cj* MlaC in *E. coli* requires periplasmic localization. The *Cj* MlaC phenotype suggests contribution to the maintenance of outer membrane integrity under antibiotic stress independent of the native *E. coli* Mla system.

### *Cj* MlaC co-purifies with endogenous phospholipids

To investigate whether *Cj* MlaC functions as a phospholipid-binding protein in *E. coli*, analogous to other MlaC homologs^32,39,40^, *Cj* MlaC was purified after expression in *E. coli* BL21(DE3) (Figure S4) and subjected to a ^31^P-NMR analysis to detect the phosphorus-containing head group of potentially associated phospholipids (Figure 2A). To specifically detect tightly bound phospholipids to purified *Cj* MlaC, membrane fractions were removed immediately after cell lysis by ultracentrifugation, and *Cj* MlaC was extensively washed while bound to the affinity matrix before elution. ^31^P-NMR analysis showed phosphorus signals that were absent in the delipidated *Cj* MlaC sample, indicating successful removal of bound phospholipids by detergent treatment (Figure 2A). To determine the lipid preference of *Cj* MlaC, phospholipids associated with the purified protein were extracted using a methanol/chloroform mixture and analyzed by TLC revealing that *Cj* MlaC co-purifies preferentially with phosphatidylethanolamine (PE) and to a lesser extent with phosphatidylglycerol (PG) (Figure 2B), consistent with the known relative abundance of these phospholipids in *E*. *coli* membranes (PE ∼75%, PG ∼20%, cardiolipin (CL) ∼5%^41^). We could not identify a clear spot for CL via TLC, most likely explained by migration of CL together with the loading front (Figure 2B). A similar lipid migration pattern was observed for lipids extracted from *Ec* MlaC, while no lipids were found associated with *Cj* MlaC after detergent-facilitated delipidation. As negative control, we used an affinity-purified sybody, not expected to tightly bind phospholipids. The fact that lipids co-purify exclusively with MlaC and not with the negative control demonstrates the specificity of phospholipid binding to MlaC (Figure 2B-C).

**Figure 2.**
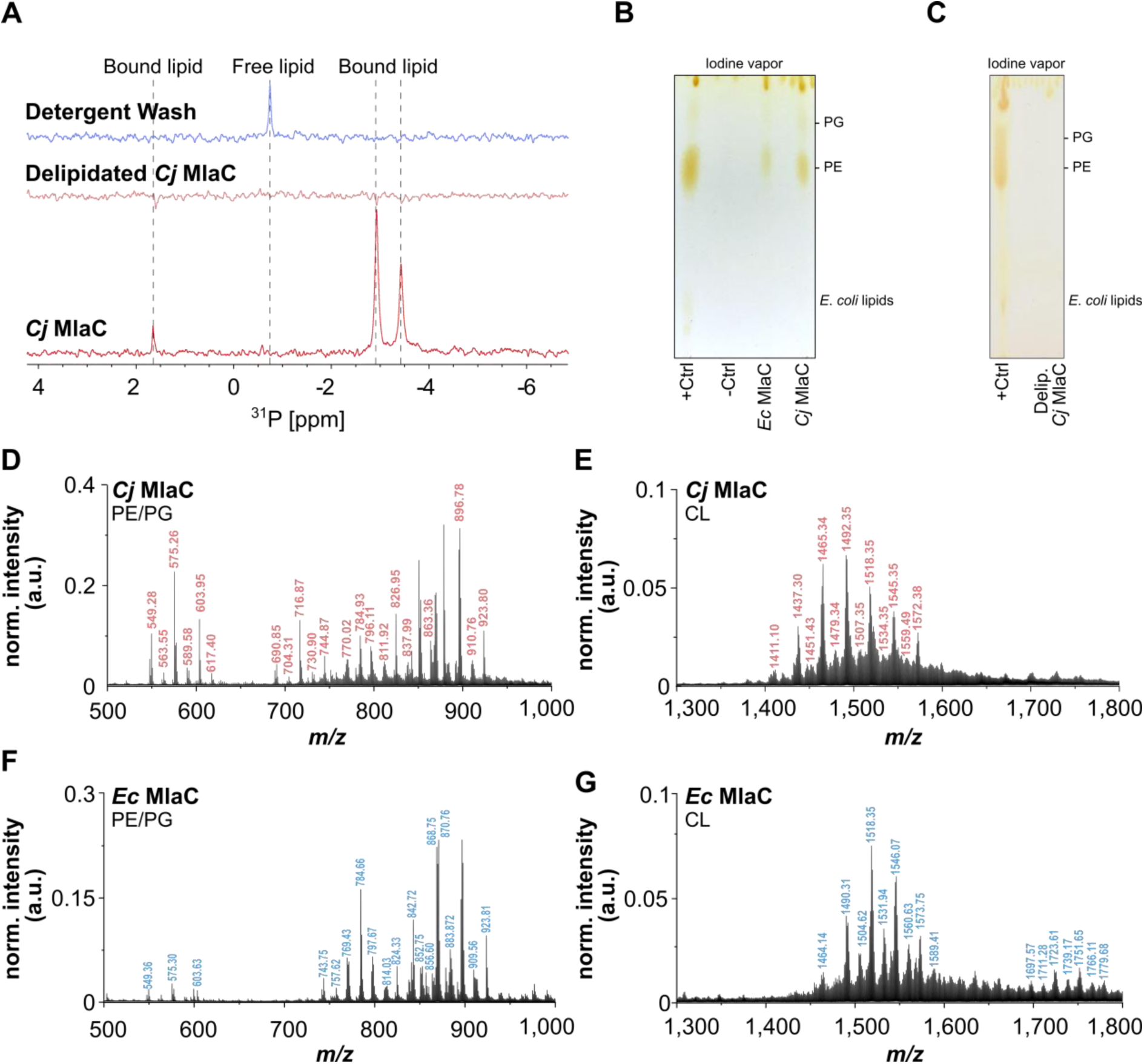
Lipid preference of MlaC. (**A**) ^31^P NMR analysis of purified *C*. *jejuni Cj* MlaC, delipidated *Cj* MlaC after incubation with the detergent *n*-nonyl-β-D-glucopyranoside (β-NG), and the detergent wash fraction to detect co-purified phospholipids. **(B)** Phospholipids co-purified with *Cj* and *E*. *coli* (*Ec*) MlaC were extracted and analyzed by thin-layer chromatography (TLC). Lipids were separated on TLC plates using a chloroform:methanol:acetic acid solvent system (65:25:10, v/v/v) and visualized by staining with iodine vapor. Total lipid extracts from *E*. *coli* served as a positive control (+Ctrl), while lipids extracted from a sybody were used as negative control (-Ctrl). The major *E*. *coli* phospholipid (PL) species were identified based on migration relative to the positive control and include phosphatidylethanolamine (PE) and phosphatidylglycerol (PG). (**C**) TLC analysis of delipidated *Cj* MlaC after incubation with β-NG for the removal of co-purified lipids. (**D-G**) Lipid extracts from purified *Cj* and *Ec* MlaC were analyzed by nano-electrospray ionization mass spectrometry (nESI-MS) to identify co-purified phospholipids. Peaks that were unambiguously assigned to one of the major *E*. *coli* phospholipid species using the MAPS® lipid database and that were consistently detected in all three biological replicates were marked and listed in Table S3. The remaining biological replicates are shown in Figure S5 and S6.

To compare the specificity profiles of the phospholipids bound to *Cj* MlaC and *Ec* MlaC, lipid extracts were analyzed by nESI-MS, showing multiple peaks in the mass ranges of 700–900 Da and 1300-1500 Da. The different phospholipid species were identified using the MAPS® lipid database (Sud *et al*., 2007; Conroy *et al*., 2024) and assigned to the major *E*. *coli* phospholipids PE, PG, CL and lyso-PLs species (Table S3). The signal intensity of the CL species was diminished, which is consistent with its lower abundance in *E. coli* (Figure 2D-G). Collectively, the results demonstrate that overexpressed *Cj* MlaC in *E*. *coli* co-purifies with phospholipids in a defined and reproducible manner. The observed specificity profile is consistent across the biological replicates and indicates that while *Cj* MlaC and *Ec* MlaC bind the same lipid classes, they have a distinct lipid preference (Figure 2D and F, Figure S6, Table S3). *Cj* MlaC co-purified with significantly more lyso-PGs than *Ec* MlaC (7.4 ± 0.6% vs. 2.4 ± 0.8% of total co-purified lipids, n = 3 biological replicates), with a consistently broader range of acyl chain lengths (19:0, 20:0, 22:1) across all replicates, compared to a single lyso-PG species (19:0) for *Ec* MlaC. Because both proteins were purified from the same *E. coli* host and therefore had access to an identical lipid pool, this discrepancy cannot be explained by differences in lyso-PL availability. This indicates an intrinsic, evolutionarily divergent binding selectivity between the two orthologs, despite both proteins binding the same underlying lipid classes.

### *Cj* MlaC forms dimers, with each monomer capable of binding phospholipids

The co-purified *Cj* MlaC-lipid complex was further analyzed via LILBID-MS to address the lipid-protein stoichiometry. Unexpectedly, *Cj* MlaC was detected in both monomeric and dimeric states, in contrast to its *E. coli* counterpart, which was observed exclusively as a monomer (Figure 3A). Furthermore, delipidated *Cj* MlaC still showed dimers, albeit to a lesser extent, suggesting an interplay between lipid presence and complex stability (Figure 3A). To determine the number of phospholipids bound by *Cj* MlaC, higher-resolution nESI-MS experiments were conducted (Figure 3B). Notably, these spectra showed additional peaks indicating a mass of approximately 800 Da bound to the monomeric form, which is missing in the spectra of delipidated *Cj* MlaC (Figure S8B). This aligns well with the typical mass range of PE or PG species. No binding of CL was detected, consistent with its low abundance in *E. coli*, which likely places it below the detection limit. The dimeric *Cj* MlaC appears to bind two of these phospholipids, one to each protomer. By increasing the collision voltage, and therefore the kinetic energy of the protein prior to impact with the inert collision gas, we were able to dissociate the bound lipids in a stepwise manner. Lipids are bound with high affinity, as high collision voltages were required to dissociate the phospholipids from both the *Cj* MlaC monomer and dimer (Figure 3B). Interestingly, at collision energies sufficient to strip most lipids of the monomeric *Cj* MlaC, the *Cj* MlaC dimer is still dominantly present with two bound lipids. At a collision voltage of 50 V, the lipid-bound fraction of the *Cj* MlaC dimer was 2-fold higher compared to the *Cj* MlaC monomer (Figure 3C). This suggests that dimerization may increase lipid affinity and therefore hints at the dimer being the preferred lipid-bound state.

**Figure 3.**
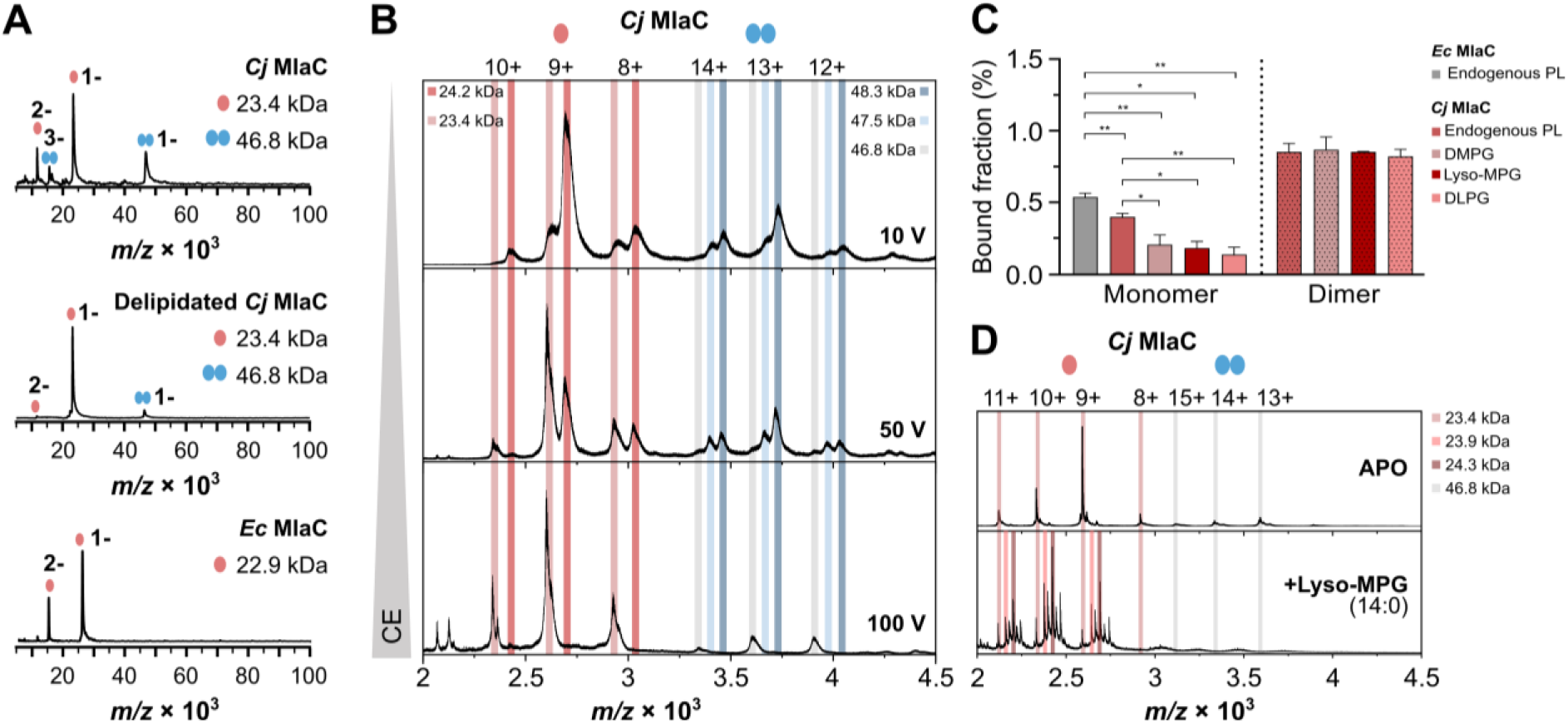
Conformational state and lipid occupancy of *C*. *jejuni* MlaC. (**A**) Laser-induced liquid bead ion desorption mass spectrometry (LILBID-MS) analysis of *C*. *jejuni* (*Cj*) MlaC, delipidated *Cj* MlaC, and *E*. *coli* (*Ec*) MlaC. Peaks were assigned to monomers (red) or dimers (blue) in different charge states. (**B**) Nano-electrospray ionization mass spectrometry (nESI-MS) analysis of *Cj* MlaC co-purified with endogenous phospholipids at different collision energies (CE). (**C**) For the *Cj* MlaC monomer and dimer, the lipid-bound peak integrals were determined for MlaC-lipid complexes after delipidation and lipid incubation or purified with endogenous lipids and compared with those of the lipid-bound *Ec* MlaC monomer. Integrals were evaluated from mass spectra acquired with a collision energy of 50 V to quantify the fraction of lipid-bound MlaC remaining after partial collision-induced activation (Table S4). Each data point represents the mean ± standard error of the mean (SEM) from three biological replicates. A two-tailed unpaired t-test was performed: * p ≤ 0.05; ** p ≤ 0.01; *** p ≤ 0.001; **** p ≤ 0.0001. (**D**) nESI-MS analysis of delipidated *Cj* MlaC in the apo form or after incubation with lyso-MPG. Peaks were assigned to the monomeric (red) or dimeric (gray) state in different charge states. The full set of replicates is shown in Figure S7 – S11.

Intriguingly, the *Ec* MlaC monomer appears to bind endogenous phospholipids much tighter than the *Cj* MlaC monomer, as evidenced by the significantly higher lipid-bound fraction in the *Ec* MlaC spectra (Figure 3C and S9A). Comparison of the collision energies required to dissociate lipids from *Cj* MlaC and *Ec* MlaC suggests that the energy required to remove lipids increases in this order: *Cj* MlaC monomer, *Ec* MlaC monomer, *Cj* MlaC dimer (Figure 3C, S8A and S9A). Collectively, these findings suggest that *Cj* MlaC and *Ec* MlaC employ distinct mechanisms for phospholipid release. This observation is further supported by the delipidation efficiencies of the *Cj* MlaC and *Ec* MlaC monomers following detergent treatment (Figure S8B and S9B). *Cj* MlaC monomers exhibited a delipidation efficiency of 89 ± 2%, compared with 47 ± 10% for *Ec* MlaC. For the *Cj* MlaC dimer, no delipidation efficiency could be determined because the dimer abundance decreased substantially following lipid removal (Figure S8B), suggesting that delipidation destabilizes the dimer and promotes its dissociation into monomers.

Next, we assessed whether *in vitro* lipid loading experiments could reveal a preference of *Cj* MlaC for specific lipids. For this, delipidated *Cj* MlaC was incubated *in vitro* with different lipid binding candidates in mixed lipid/β-NG micelles and subsequently analyzed by nESI-MS. Following incubation of delipidated *Cj* MlaC with the micelles (1:5 protein:lipid molar ratio), we could observe binding of phospholipids with a PG headgroup, while neither PE nor CL phospholipid species showed detectable interaction with *Cj* MlaC (Figure S10A). Furthermore, incubation of *Cj* MlaC with PG species differing in acyl chain length revealed binding of a single phospholipid to each protomer of the *Cj* MlaC monomer and dimer, with lipid-loading efficiency increasing as acyl chain length decreased. The largest number of bound complexes was observed for DLPG, followed by DMPG (Figure S10A and S11). Because the observed differences in lipid-loading efficiency as a function of acyl chain length could reflect enhanced accessibility of short-chain lipids to *Cj* MlaC or higher binding affinity, we compared the lipid-bound fraction of *Cj* MlaC after increasing the collision voltage to 50 V to assess protein-lipid complex stability. The amount of the *Cj* MlaC dimer that retained bound lipids at 50 V was approximately the same for DMPG, lyso-MPG, DLPG, and for the endogenous lipid. In contrast the fraction of lipid-bound *Cj* MlaC monomer differed considerably between the phospholipid species (Figure 3C). While *Cj* MlaC retained fewer endogenous lipids than the *Ec* MlaC, the proportion of bound DMPG and DLPG in monomeric *Cj* MlaC was markedly lower than that of the endogenous lipid, indicating an increased affinity for the endogenous, longer chain-length phospholipid. Interestingly, the lipid affinity appeared to increase upon protein dimerization, independently of acyl chain length, confirming the earlier observation that lipid affinity is higher for dimeric *Cj* Mlac (Figure 3C and S8A).

So far, we have observed binding of one lipid per protomer. For probing the binding stoichiometry between MlaC and the number of acyl chains, *Cj* MlaC was lipidated *in vitro* with lyso-MPG, a monoacyl PG species with a 14-carbon acyl chain (Figure 3D). The mass spectra showed binding of one or two lyso-MPG molecules per *Cj* MlaC protomer. Unexpectedly, three additional lipid-bound *Cj* MlaC species were observed (Figure 3D and S10B). These species may result from the binding of a lyso-MPG molecule that is 156.2 Da heavier, potentially corresponding to a 10-carbon acyl chain moiety and probably derived from a synthetic intermediate formed during lipid synthesis or from modifications occurring during storage of the lyso-MPG sample. Across all lyso-MPG species, we detected a maximum of two bound molecules per MlaC protomer, reinforcing the notion that *Cj* MlaC can accommodate one or two acyl chains of a phospholipid within its lipid binding pocket. This agrees with our previous experiments showing consistent binding of only one diacyl phospholipid (Figure 3B).

### *Cj* MlaC first undergoes a conformational change prior to phospholipid release

To gain further insights into the lipid binding mechanism of *Cj* MlaC, we conducted a combination of Tandem MS/MS and collision induced unfolding ion mobility mass spectrometry (CIU IMS). In the MS/MS experiments of *Cj* MlaC bound to its endogenous, co-purified lipids, the 13+ charge state of the dimer bound to two phospholipids was mass-selected and transferred into the collision cell for controlled activation. A stepwise increase in the collision energy resulted in the sequential dissociation of the lipids at collision energies above 50 V (Figure 4A). At 80 V, almost no dimer with two bound lipids remained, and the spectrum was dominated by lipid-free *Cj* MlaC dimer. Only after both lipids were dissociated from the *Cj* MlaC dimer, a further increase in energy led to the dissociation of the now lipid-free dimer into its monomeric state (Figure 4A). Intriguingly, we did not observe any phospholipid-bound monomer as a result of collision-induced dissociation (CID), suggesting that the interaction between the two *Cj* MlaC monomers is stronger than that between each protomer and the phospholipid. Moreover, when the MS/MS experiments were repeated by selecting the lipid-bound *Cj* MlaC monomer (Figure 4B), complete lipid dissociation was achieved at a lower collision energy of 70 V (for the 8+ charge state, correlating to a lab frame energy of 560 eV) in contrast to the dimer, which required collision voltages above 100 V (charge state 13+, lab frame energies exceeding 1300 eV), again hinting at higher lipid binding affinity of the dimer.

**Figure 4.**
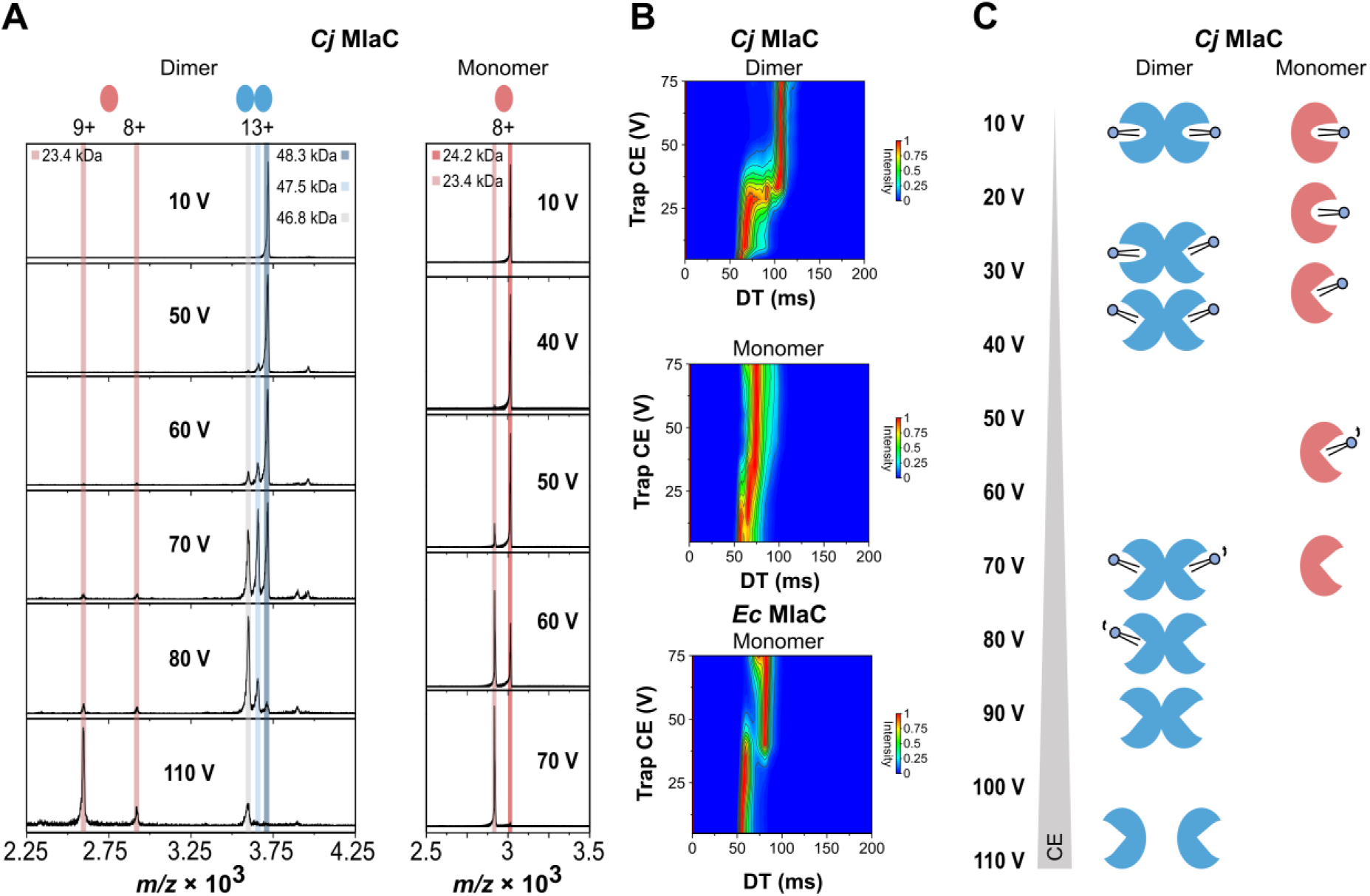
Conformational dynamics of *Cj* MlaC during lipid and dimer dissociation. (**A**) Tandem mass spectrometry (MS/MS) analysis *of Cj* MlaC bound to its endogenous, co-purified lipids at different collision voltages filtered for masses corresponding to the *Cj* MlaC dimer or monomer. (**B**) Collision-induced unfolding ion mobility spectrometry (CIU IMS) analysis of a lipid-bound *Cj* MlaC dimer (13+), monomer peak (8+) or lipid-bound *Ec* MlaC monomer peak (8+). The trap collision energy (CE) was plotted as a function of the drift time (DT). The full set of replicates is shown in Figure S12 and S13. (**C**) Proposed phospholipid dissociation pathway of *Cj* MlaC based on our *in vitro* experiments, illustrating lipid and dimer dissociation for *Cj* MlaC. *Cj* MlaC can exist as monomer and dimer. Application of high collision voltages leads to a structural opening of each *Cj* MlaC protomer, likely expanding the lipid-binding pocket and thus facilitating the release of the bound phospholipid. The structural rearrangement of the dimer is larger than for the monomer and requires higher energies. Further increase in collision voltage results in dissociation of the now lipid-free dimer into monomers, suggesting that the affinity of *Cj* MlaC for phospholipid is weaker than for a second *Cj* MlaC monomer to form dimers.

The CIU IMS data for the *Cj* MlaC dimer furthermore revealed that the dimer undergoes a stepwise structural rearrangement, with either two distinct transitions very close in energy (35 V and 40 V, Figure 4B and S12C), or both occurring at the same energy. These structural rearrangements occur at collision energies just below those for which the MS/MS data indicates the dissociation of the bound phospholipids in a stepwise manner (50 V) (Figure 4A). This supports the interpretation that the observed unfolding transitions correspond to the opening of the lipid-binding cavities and that the bound phospholipids are tightly enclosed within the lipid-binding cavity. Remarkably, the IMS spectra of the lipid-free *Cj* MlaC dimer showed a comparable structural rearrangement at similar collision energies (Figure S12E). This might suggest that, in the lipid-free state, the lipid-binding pocket of a *Cj* MlaC dimer adopts a similar closed conformation. The Tandem MS/MS and CIU IMS data for the monomeric state showed likewise a similar behavior, though the effect was less pronounced (Figure 4A-B). Thereby, monomeric *Cj* MlaC exhibited a small first unfolding step followed by a more gradual drift time shift between 15 V and 35 V, suggesting likewise a structural rearrangement of the protein prior to releasing its single bound phospholipid at a collision energy from around 40 V (Figure 4A-B). The lack of the two larger unfolding events, observed for the dimer suggests the opening of the single lipid-binding cavity in the monomer. The more gradual unfolding indicates a less rigid structural element. Remarkably, the CIU IMS data for the *Ec* MlaC homolog showed a considerably larger and a more defined drift time shift than for the *Cj* MlaC monomer (Figure 4B), increasing from approximately 55 ms to 80 ms. This shift results from a more pronounced unfolding event, showing greater similarity to the *Cj* MlaC dimer than to the *Cj* MlaC monomer. This could indicate that the *Ec* MlaC monomer, like the *Cj* MlaC dimer, adopts a more defined compact conformation allowing for a tighter association with its bound lipids, while the *Cj* MlaC monomer exists in a more loosely packed state. This is consistent with our previous findings showing lower delipidation efficiency of *Ec* MlaC by detergent treatment and a higher lipid-bound fraction of the *Ec* MlaC monomer compared to the *Cj* MlaC monomer despite increasing the collision energies (Figure 3C and S9).

Altogether, these findings support a model in which *Cj* MlaC takes on a more loosely packed form as a monomer, whereas lipid-loading promotes the dimeric form and dimerization stabilizes the lipid-bound state. Subsequent structural rearrangement is required to allow for stepwise phospholipid release from each monomer within the *Cj* MlaC dimer, followed by lipid-free dimer dissociation into two monomers.

## Discussion

The *E. coli* Mla pathway plays a critical role in maintaining OM asymmetry by transporting mislocalized phospholipids from the outer leaflet of the OM to the inner membrane^6^. The OM-localized MlaA thereby catalyzes the transfer of phospholipids from the outer leaflet onto the periplasmic phospholipid shuttle MlaC. Phospholipid-loaded MlaC then diffuses through the periplasm and contacts the MlaD ring of the IM-localized MlaEFDB ABC transporter. After phospholipid transfer, the phospholipids are incorporated into the IM lipid bilayer upon ATP hydrolysis. Differently, in *C*. *jejuni*, the genes encoding the ABC transporter complex MlaEFD are encoded on a different operon than the genes for the lipid transfer proteins MlaA (*Cj*1371) and MlaC (*Cj*1372), which are co-transcribed with a gene encoding an RND transporter (*Cj*1373) (Figure S1A). Interestingly, the genes for MlaC and the RND transporter overlap with their stop and start codon (UAAUG), respectively, indicating that these genes are translationally coupled and therefore that their synthesis is tightly associated. Given that *C. jejuni* encodes the Mla system together with an RND transporter, whose members usually function as substrate/H⁺ antiporters, we hypothesize that the *C. jejuni* Mla system may serve dual functions: mediating (1) retrograde phospholipid transport via MlaA and MlaFDB and (2) anterograde transport via *Cj* MlaA, MlaC, and the same-operon-encoded RND antiporter (Figure S1B). The aim of this study was to gain functional and mechanistic insights into the role of the periplasmic *Cj* MlaC protein in phospholipid transport.

In this study, functional characterization of *Cj* MlaC in different *E. coli* BW25113 *mla*-deficient strains revealed that *Cj* MlaC enhances the cell proliferation of all mutant strains under antibiotic stress. By contrast, a cytosolically expressed *Cj* MlaC variant resulted in little or no growth enhancement, indicating that the periplasmic localization is essential for its function (Figure 1). Even when the cells were exposed to vancomycin, a large antibiotic that cannot bind to MlaC due to size restriction and that cannot readily cross the outer membrane unless compromised, the same growth-promoting effect of *Cj* MlaC was perceived, emphasizing that this effect is not due to direct antibiotic sequestration via *Cj* MlaC but rather hinted toward the restoration of the OM asymmetry (Figure S2B). However, in contrast to *Ec* MlaC, which can only complement an *E. coli mlaC*-deficient strain, *Cj* MlaC enhanced cell growth in the presence of antibiotics regardless of the specific deletion background. Intriguingly, this effect was most pronounced in an *mlaE*-deficient strain, which serves as component of the energy-driven ABC transporter of the Mla system, demonstrating a strong growth-promoting role of *Cj* MlaC in *E. coli*. These findings suggest that *Cj* MlaC functions independently of the endogenous *E. coli* Mla pathway and may instead bind and release phospholipids independently, possibly also in both directions in *E. coli*. Moreover, this study revealed that *Cj* MlaC co-purifies with phospholipids after extensive wash steps (Figure 2), suggesting that it has a high affinity for phospholipids and confirms that it functions as a phospholipid-binding protein. Determination of the bound phospholipid species to purified *Cj* MlaC revealed PE as the most abundant co-purified lipid species, followed by PG, and to a much lesser extent CL (Figure 2B, D-E). This is consistent with the relative abundances of these phospholipid species in *E*. *coli*^41^ and with the results obtained for *Ec* MlaC (Figure 2B, F-G)^32^. Interestingly, while *Cj* and *Ec* MlaC bind the same lipid classes, they exhibit distinct lipid-binding preferences^42^, as reflected by the differing *m/z* values in the nESI-MS spectra of the extracted bound lipids, corresponding to variations in acyl-chain lengths and saturation (Figure 2D-G, Figure S6, Table S3). Remarkably, *Cj* MlaC co-purified with significantly more lyso-PG than *Ec* MlaC (7.4 ± 0.6% vs. 2.4 ± 0.8% of total co-purified lipids, n = 3 biological replicates; Table S5), spanning a broader range of acyl chain lengths (19:0, 20:0, 22:1) versus a single species (19:0) for *Ec* MlaC. Since both proteins were purified from the same *E. coli* host and thus sampled an identical lipid pool, this divergence cannot reflect differences in lyso-PL availability but instead points to an intrinsic difference in binding-pocket selectivity between the two orthologs.

This is consistent with the distinct physiological demands placed on the Mla pathway in each organism. Lyso-PLs are a usually particularly destabilizing lipid species, and *C*. *jejuni* lacks the LplT-Aas repair system that *E*. *coli* uses to rapidly clear them, allowing lyso-PLs to accumulate to 28–45% of total membrane lipids, compared to <1% in most other Gram-negative bacteria^21^. The elevated and more acyl chain length-diverse lyso-PG binding of *Cj* MlaC may reflect a binding pocket adapted for retrieval of these lipids. This raises the possibility that *Cj* MlaC might be involved in RND-coupled anterograde delivery of lyso-PLs to the outer membrane, where it has been reported that the presence of these lipids are correlated with *C. jejuni* motility, a property crucially important to colonize the cecum of chicken^22^.

Intriguingly, we observed that purified *Cj* MlaC can exist in both monomeric and dimeric states, unlike *Ec* MlaC and other homologs, which have been reported exclusively as monomers (Figure 3A)^8,39^. In our nESI-MS analysis with purified *Cj* MlaC, each protomer was consistently observed to bind a mass of approximately 800 Da (Figure 3B), which falls within the mass range of common PE or PG molecular species suggesting the binding of a single phospholipid per protomer. Moreover, CID experiments showed that lipids can be dissociated from both the monomer or dimer. Interestingly, the dissociating phospholipids from the monomeric *Cj* MlaC or the *Ec* MlaC required much lower collision energies than for those bound to the dimeric *Cj* MlaC, indicating that dimerization increases lipid-binding affinity. This correlates well with the delipidation efficiencies determined for *Ec* MlaC and *Cj* MlaC, showing that detergent treatment removes lipids from the *Ec* MlaC monomer only half as efficiently as from the *Cj* MlaC monomer, which concurrently causes dissociation of the *Cj* MlaC dimer (Figure S8B, S9B and Table S4). These findings show that the unique dimerization of *Cj* MlaC causes higher lipid affinity, and the dimer is concomitantly stabilized. As a next step, we tested the feasibility of lipid delivery *in vitro*, showing that mixed lipid/β-NG micelles are suitable vehicles for incubation studies with lipid binding proteins. *In vivo* binding behavior, inferred from the identification of phospholipids co-purified with *Cj* MlaC, indicated binding of a variety of PE, PG, and CL species (Figure 2D-E, Table S3). This observation points toward a headgroup-independent specificity as shown before for *Ec* MlaC which has been reported to primarily recognize the acyl chains of the phospholipids bound^43^. Interestingly, in the *in vitro* re-lipidation experiment, we found that only PG lipid species were successfully delivered from ß-NG micelles to bind to *Cj* MlaC. The preferential binding of PG lipids to *Cj* MlaC monomer and dimer may be best explained by their higher accessibility within the micelles compared to PE and CL micelles^43^. Therefore, we conducted our subsequent analysis of the similarities and differences in the lipid-binding mechanisms of *Ec* MlaC and *Cj* Mlac using PG lipids as representative ligands. Lipid loading was particularly efficient with decreasing acyl chain length, which we interpret as a consequence of the mixed β-NG/short-chain lipid micelle properties that render these lipids more accessible to *Cj* MlaC and does not necessarily support the notion of stronger binding.

Recognizing the need for deeper mechanistic insight into lipid binding, we compared the protein-lipid complex stabilities across different bound lipid species at a collision energy of 50 V (Figure 3C). Intriguingly, shorter chain DMPG- and DLPG-lipids were released more readily from *Cj* MlaC monomers than the longer acyl chain endogenous phospholipids, which on average contain acyl chains of at least 16 carbon atoms^44^. This acyl chain length dependence was not observed for the *Cj* MlaC dimer, which as well retained the lipids much more strongly than the monomer. In addition, our *in vitro* lipidation studies with lyso-MPG suggest that a *Cj* MlaC monomer can simultaneously engage both acyl chains of two monoacyl PLs or of one diacyl PL.

To further investigate the differences in lipid binding, we sought to gain deeper insight into the architecture of the binding cavities, as structural features might account for the observed variations. Comparison of the collision energies required for lipid release with our CIU IMS data showed that the *Cj* MlaC monomer and dimer have to adopt an extended conformation prior to lipid release (Figure 4A-B). This effect is more pronounced in the dimeric form, where the *Cj* MlaC dimer is observed to undergo two concerted unfolding events in quick succession, probably reflecting an opening event of each protomer. In comparison, the *Cj* MlaC monomer showed only a small drift-time shift already at low collision energies, followed by a gradual increase across the activation range, suggesting a less well-defined, less stable structure than for the dimer. Furthermore, we observed that the structural rearrangement of the *Ec* MlaC monomer was much more pronounced than that of the *Cj* MlaC monomer, requiring higher collision energies and exhibiting unfolding features more comparable to the *Cj* MlaC dimer than to the monomer. These larger conformational changes might reflect a

transition from a distinct closed to an open state, explaining the higher phospholipid-binding affinity of *Ec* MlaC and *Cj* MlaC dimer. In contrast, the more loosely packed structure of monomeric *Cj* MlaC is indicative of a less rigid overall architecture. This apparent flexibility correlates with the enhanced capacity of *Cj* MlaC to bind lyso-PLs, as these single-chain lipids are sterically less demanding and may not require as tightly closed a pocket conformation as diacylated species for initial binding. Moreover, our CIU IMS data for the lipid-free *Cj* MlaC dimer showed that it exhibits a similar unfolding behavior as the lipid-bound dimer (Figure S12E). This might suggest that the lipid-free *Cj* MlaC dimer adopts a closed conformation, analogous to the closed state previously described for monomeric *Ec* MlaC^45^.

Tandem MS/MS analysis of *Cj* MlaC bound to its endogenous and co-purified lipids as well, revealed sequential lipid release from the *Cj* MlaC dimer prior to dimer dissociation, indicating a stronger interaction between the two *Cj* MlaC protomers than of a protomer to a phospholipid (Figure 4A). Notably, we observed that the *Ec* MlaC monomer was less efficiently delipidated by detergent than the *Cj* MlaC monomer and dimer (Figure S8B and S9B, Table S4), even though we observed that the phospholipids appear to be more tightly associated with the *Cj* MlaC dimer (Figure 3C). The differences in delipidation efficiencies between detergent-mediated delipidation and collision-induced phospholipid release in the MS experiments might be attributable to the longer exposure to detergent (overnight) and a potential cooperative mechanism, in which the release of one phospholipid destabilizes binding of the second phospholipid in the opposing protomer, ultimately leading to dissociation of the dimer into two monomers.

Combining all our findings from the *in vitro* lipidation studies, the CIU IMS data and the tandem MS/MS analysis, we can conclude that while lipids with shorter acyl chains are more readily bound by *Cj* MlaC in our binding assays, this behavior does not translate into stronger binding. Even mild activation at a collision energy of 50 V led to lipid dissociation from the *Cj* MlaC monomer in a manner that correlates with acyl chain length, with shorter acyl chain lipids dissociating more readily (Figure 3C). In comparison, *Cj* MlaC dimers bind their lipids even more strongly, independent of the acyl-chain length (Figure 3C). A comparison of the tandem MS/MS and the respective CIU energy profiles indicate that in all cases lipid release is coupled to protein conformational change and proceeds only after the protein adopts an open state. Nevertheless, the CIU patterns show very informative differences (Figure 4B). The *Cj* MlaC monomer adopts a loosely packed, intrinsically unstable conformation that begins to gradually expand upon activation already at low collision energies. This suggests a more flexible structure, which might support the enhanced binding of lyso-PLs, as these single-chain lipids are sterically less demanding and may not require as tightly closed a pocket conformation as diacylated species. In such a configuration, where no lid-like structure retains the lipids, the extent of the contact surface between lipid and protein appears to determine lipid retention: lipids with shorter acyl chains, such as the 12-carbon DLPG, exhibit a reduced contact area and dissociate more readily. This behavior contrasts sharply with the CIU profile of *Ec* MlaC, which displays a more compact and structurally well-defined state, as well as a distinct opening step in the CIU plot leading at higher collision energies to a clearly wider open conformation prior to lipid loss (Figure 4B). Notably, the weak lipid-binding characteristics of monomeric *Cj* MlaC are mitigated in the dimeric form, which exhibits more defined CIU patterns and thereby resembles the structural behavior of *Ec* MlaC (Figure 4B). In this structurally well-defined dimeric state, which shows a clear distinction between the lipid-retaining closed conformation and the lipid-releasing open conformation, the *Cj* MlaC dimer retains lipids markedly more effectively, independent of lipid size. Based on the findings of this study, we propose that in *C. jejuni*, lipid-free *Cj* MlaC initially adopts a relatively loosely packed closed, low-lipid-affinity monomeric conformation *in vivo,* which is supportive of lipid loading, with an increased adaptation to lyso-PLs. Efficient lipid-loading from lipids out of the IM to the *Cj* MlaC monomer may be mediated by the putative lipid/H^+^ RND antiporter under consumption of the proton motive force (Figure 5). Interaction of *Cj* MlaC with the RND antiporter might thereby induce the opening of the lipid-binding cavity of *Cj* MlaC to facilitate phospholipid transfer. After lipid binding, two *Cj* MlaC monomers dimerize, resulting in a high-affinity lipid-loaded state that is structurally more rigid and prevents spontaneous lipid release within the periplasm or back to the inner membrane. After reaching the outer membrane, the *Cj* MlaC dimer may require interaction with an additional protein, such as *Cj* MlaA, to trigger a conformational transition and opening of the lipid-binding cavity, thereby enabling phospholipid release. The phospholipid release might be coupled to the *Cj* MlaC dimer dissociation into two monomers, allowing the lipid-free *Cj* MlaC monomers to re-enter a new phospholipid transfer cycle (Figure 5).

**Figure 5.**
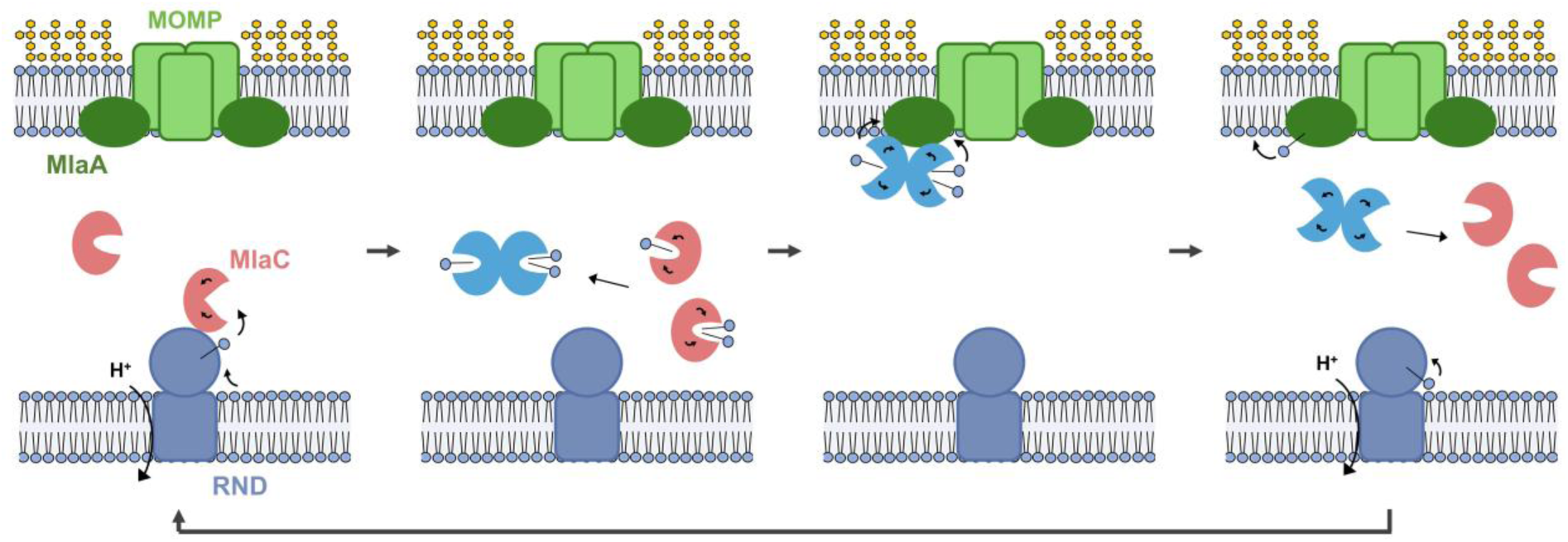
Proposed mode of action of MlaA, MlaC and the same-operon-encoded RND transporter from *C. jejuni*. Based on our findings, we propose that lipid-free *Cj* MlaC initially adopts a closed, low-lipid-affinity monomeric conformation. Lipid-free *Cj* MlaC likely interacts with the same-operon-encoded RND transporter, which is thought to energize phospholipid translocation from the inner membrane via proton antiport. This interaction might induce the opening of the phospholipid-binding pocket of *Cj* MlaC, allowing lipid binding. Lipid binding thereby induces the dimerization of *Cj* MlaC, resulting in a high-affinity lipid-loaded state. After reaching the outer membrane, the *Cj* MlaC dimer may require interaction with an additional protein, such as *Cj* MlaA, a lipoprotein potentially associated with the major outer membrane porin MOMP of *C. jejuni*, to trigger a conformational transition and opening of the lipid-binding cavity. After phospholipid release in a stepwise manner and dissociation of the dimer, the two monomers adopt again a closed, lipid-free state and are available for a new phospholipid transfer cycle. The figure emphasizes anterograde lysophospholipid transport, in line with the lysophospholipid-binding propensity of *Cj* MlaC and their prevalence in the *C. jejuni* lipidome.

In summary, our study establishes *Cj* MlaC as a phospholipid-binding protein that operates in *E. coli* independently from the native *E. coli* Mla pathway. *Cj* MlaC exhibits a unique dimerization behavior that enables transitions between a low-lipid-affinity monomeric state, with an elevated affinity for lyso-PLs compared to *Ec* MlaC and a unique high-lipid-affinity dimeric state, while accommodating no more than two acyl chains within its lipid binding pocket. This study has provided new mechanistic insights into the lipid binding mechanism of *Cj* MlaC, which will be the basis for further studies to elucidate the structural basis of *Cj* MlaC dimerization, the identity of its interaction partners, and the lipid transport pathway mediated by *Cj* MlaC with the same-operon-derived *Cj* MlaA and putative lipid/H^+^ RND antiporter in *C. jejuni.* Collectively, our results support the hypothesis that *Cj* MlaC and *Ec* MlaC fulfill distinct physiological roles *in vivo*, with *Cj* MlaC potentially contributing to anterograde phospholipid/lysophospholipid transport, optimized for the distinctive lipidome of *C. jejuni*, which maintains the membrane fluidity necessary for motility, whereas *Ec* MlaC is established to function in retrograde transport.

## Author contributions

Conceptualization: LF, TR, NM, KMP

Methodology: LF, TR, SS, LL, CB, NMB ML, WEF, HKT, AH

Investigation: LF, TR, SS, LL, CB, NMB, ML, WEF, HKT, AH

Visualization: LF, TR, CB

Supervision: NM, KMP, CG

Writing - original draft: LF, TR, NM, KMP

Writing - review and editing: LF, TR, NM, KMP

## Acknowledgements

We like to thank Paula Holzhütter (Dötsch lab, Goethe University Frankfurt) for the selection of the anti-*Cj* MlaC sybody. CG, KMP, NM acknowledge funding by the Deutsche Forschungsgemeinschaft (DFG, German Research Foundation) under Germanýs Excellence Strategy – EXC-3094 – 533751785 and SFB1507/P01, P03, P13.

## Competing interests

The authors declare that they have no competing interests.

## Notes

### Competing Interest Statement

The authors have declared no competing interest.

