## Supplemental data for "A Monomer-Dimer Equilibrium Tunes Phospholipid Handling by *Campylobacter jejuni* MlaC to the Bacterium’s Unique Lipidome"

### Supplementary figures

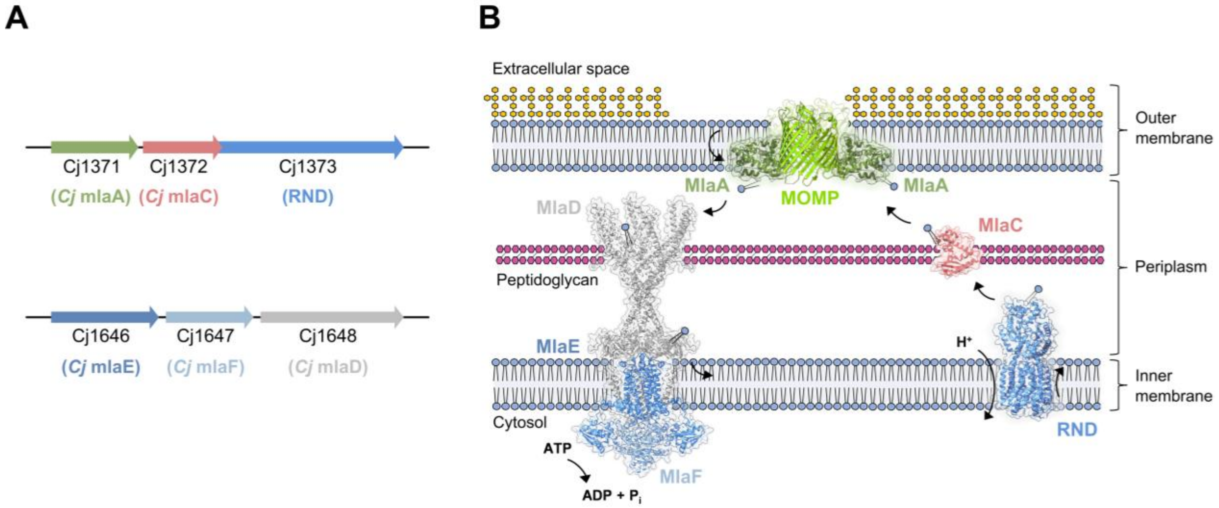


**Figure S1 | *C. jejuni* Mla system.**

(**A**) Genomic organization of the *C. jejuni* (*Cj*) Mla pathway. The genes encoding *Cj* MlaA and MlaC share an operon with an RND transporter, with the latter two being translationally fused, whereas MlaE, MlaF, and MlaD are encoded in a separate operon. (**B**) Proposed dual role of the *C. jejuni* Mla system in phospholipid transport. In retrograde phospholipid transport, the outer membrane lipoprotein MlaA (AF-Q0P8N8-F1), possibly associated with the major outer membrane porin MOMP (PDB ID: 5LDV), captures mislocalized phospholipids from the outer leaflet of the outer membrane and transfers them to the ABC transporter complex MlaEFD (*Cj* MlaE: AF-Q0P7Y2-F1, *Cj* MlaF: AF-Q0P7Y1-F1. *Cj* MlaD: AF-Q0P7Y0-F1). In anterograde phospholipid transport, the RND transporter (AF-Q0P8N6-F1) is thought to deliver newly synthesized phospholipids from the inner membrane to the periplasmic shuttle protein MlaC (AF-Q0P8N7-F1), and then to MlaA, for incorporation into the outer membrane. MlaA may function in both directions or exclusively in one, with the other pathway potentially using an alternative outer membrane component. Protein structures were visualized using UCSF ChimeraX[46].

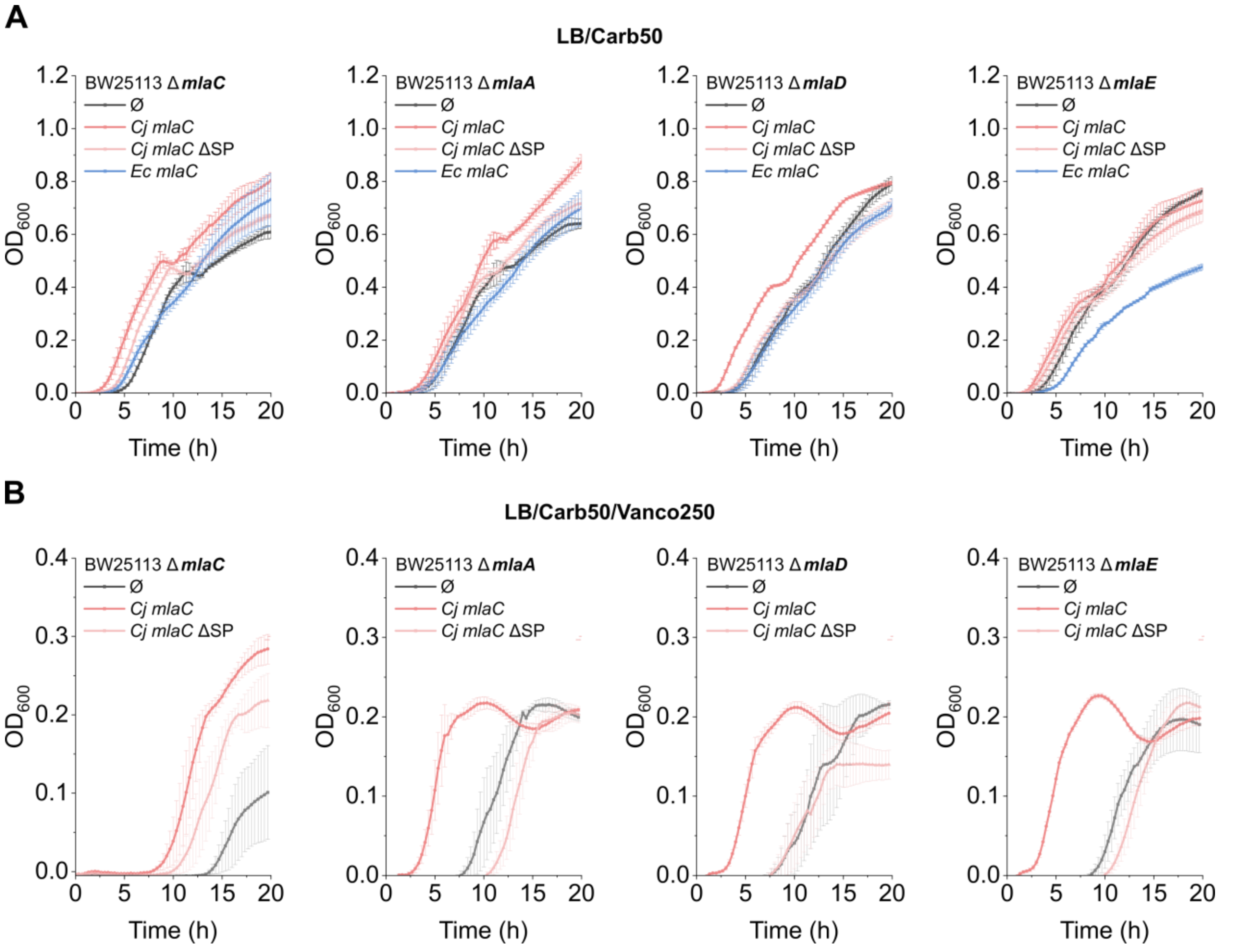


**Figure S2 | Functional characterization of *Cj* MlaC in *E*. *coli*.**

(**A-B**) Growth curve analysis of *E*. *coli* (*Ec*) BW25113 cells deficient in endogenous *mlaC*, *mlaA*, *mlaD*, or *mlaE*, and complemented with either the empty pTTQ18 vector (gray, Ø), pTTQ18 encoding *C. jejuni* (*Cj*) MlaC (dark red), the signal-peptide cleaved *Cj* MlaC variant (light red, *Cj mlaC* ΔSP) or *Ec* MlaC (blue). Cells were grown in lysogeny broth (LB) medium containing 50 µg/mL carbenicillin as selection antibiotic alone (**A**) or (**B**) with 250 µg/mL vancomycin as additional antimicrobial agent for 20 h at 37 °C. Optical density at 600 nm ($OD_{600}$) was measured at the indicated time points and plotted over time. Each data point is presented as the mean ± standard error of the mean (SEM) from three biological replicates.

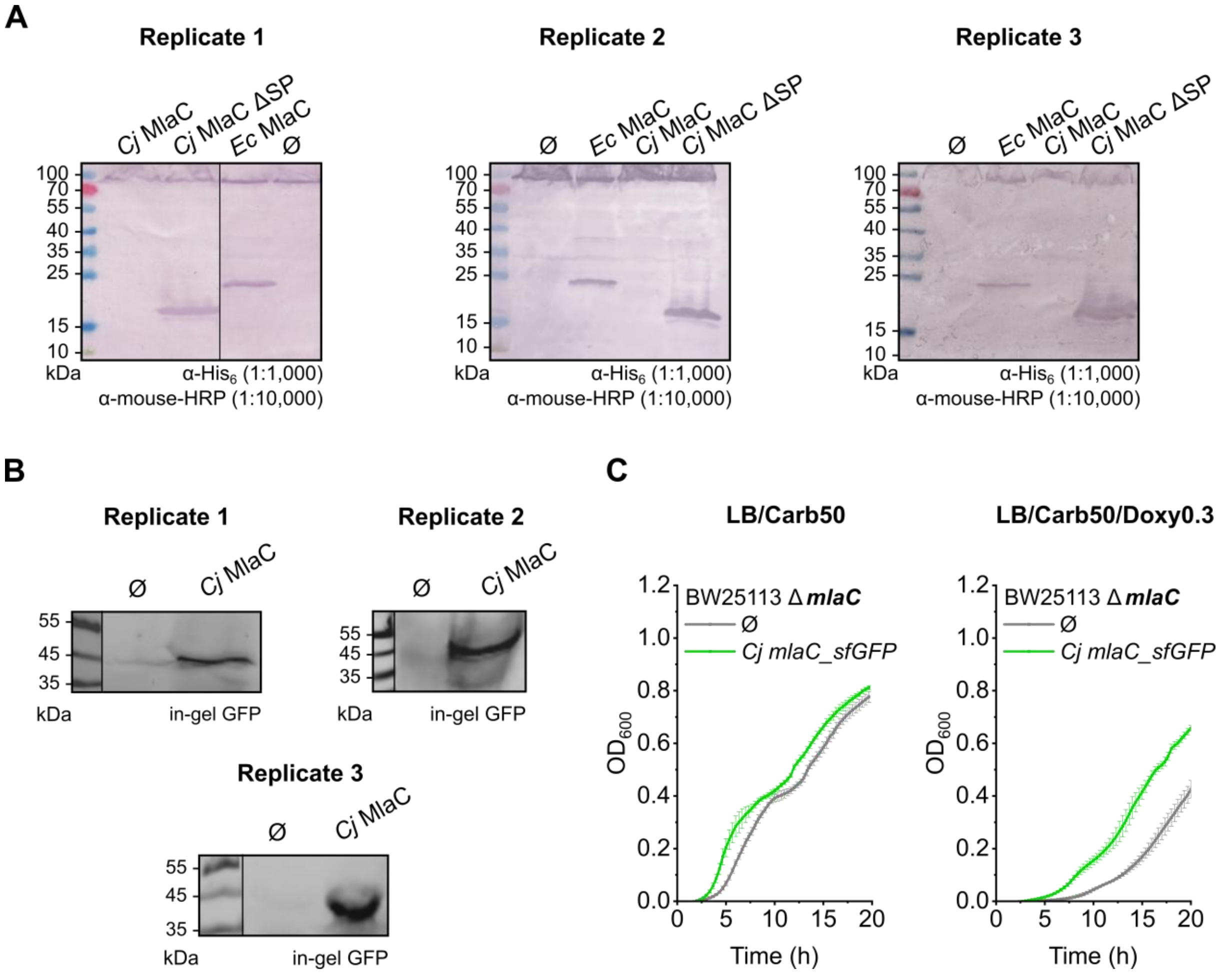


**Figure S3 | Expression analysis of pTTQ18 constructs used for functional analysis in *E. coli*.**

(**A**) Western blot analysis of the constructs used for the Growth curve analysis of MlaC in *E. coli* (*Ec*) BW25113 wild-type cells in biological triplicates. (**B**) In-gel GFP fluorescence analysis of cell lysates expressing *Cj* MlaC with a C-terminal superfolder green fluorescent protein (sfGFP) tag. All genes were expressed from the pTTQ18 vector in *E. coli* BW25113 cells. As a negative control (Ø), cell lysates from cells transformed with the empty pTTQ18 vector were applied to the gel. For each sample, 20 µL of cell lysate harvested with an $OD_{600}$ of 20 were loaded onto the gels. The Western blots and gels were cropped as indicated by the black vertical lines. (**C**) Growth curve analysis of *E. coli* BW25113 Δ*mlaC* cells complemented with either the empty pTTQ18 vector (gray, Ø) or pTTQ18 encoding *C. jejuni* (*Cj*) MlaC with a C-terminal sfGFP tag (green). Cells were grown in lysogeny broth (LB) medium containing 50 µg/mL carbenicillin (left panel) and additionally 0.3 µg/mL doxycycline (right panel) for 20 h at 37 °C. Optical density at 600 nm ($OD_{600}$) was measured at the indicated time points and plotted over time. Each data point is presented as the mean ± standard error of the mean (SEM) from three biological replicates.

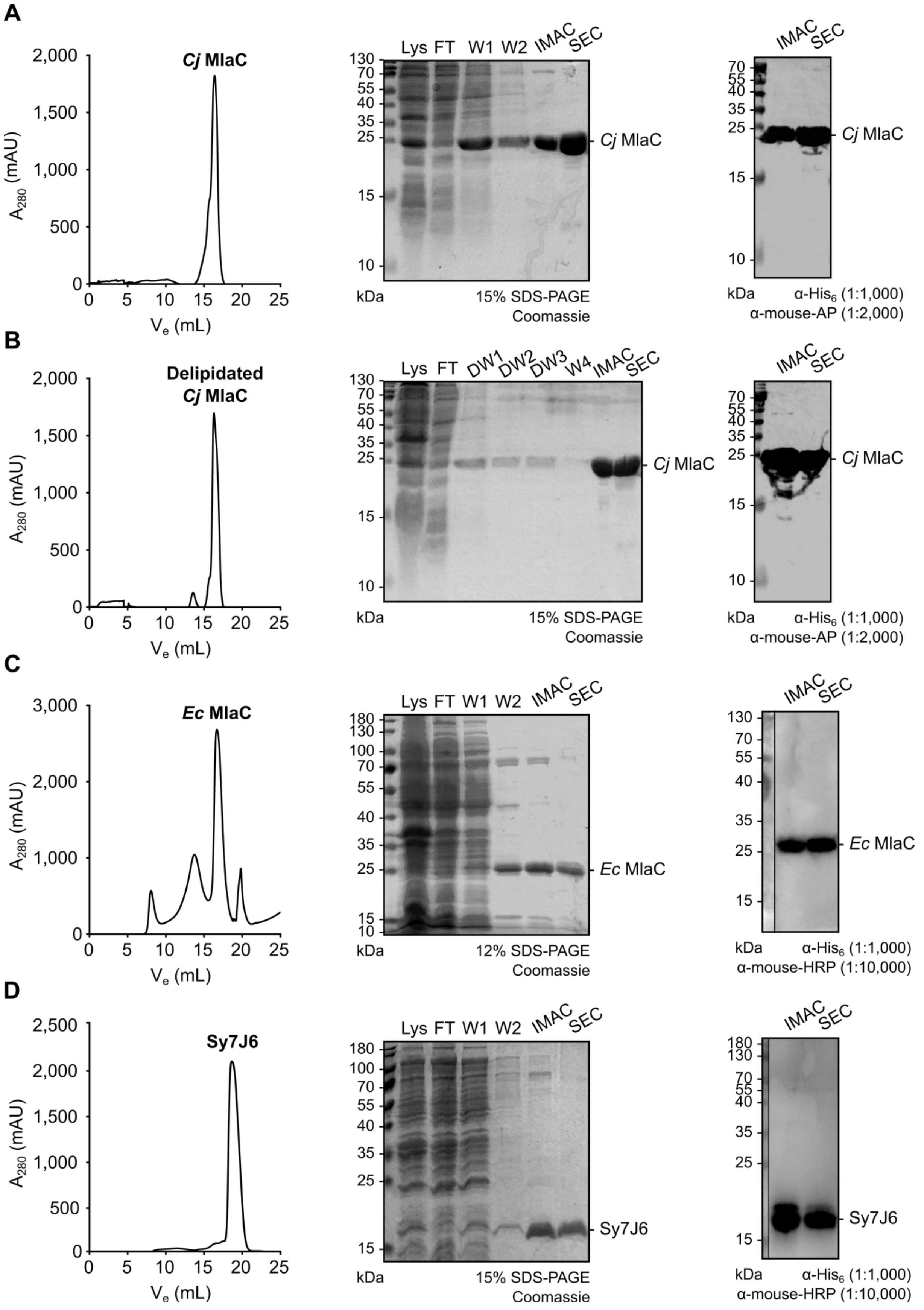
A
Cj MlaC
A280 (mAU)
Ve (mL)
Lys FT W1 W2 IMAC SEC
Cj MlaC
kDa
15% SDS-PAGE
Coomassie
IMAC SEC
Cj MlaC
kDa
α-His6 (1:1,000)
α-mouse-AP (1:2,000)
B
Delipidated
Cj MlaC
A280 (mAU)
Ve (mL)
Lys FT DW1 DW2 DW3 W4 IMAC SEC
Cj MlaC
kDa
15% SDS-PAGE
Coomassie
IMAC SEC
Cj MlaC
kDa
α-His6 (1:1,000)
α-mouse-AP (1:2,000)
C
Ec MlaC
A280 (mAU)
Ve (mL)
Lys FT W1 W2 IMAC SEC
Ec MlaC
kDa
12% SDS-PAGE
Coomassie
IMAC SEC
Ec MlaC
kDa
α-His6 (1:1,000)
α-mouse-HRP (1:10,000)
D
Sy7J6
A280 (mAU)
Ve (mL)
Lys FT W1 W2 IMAC SEC
Sy7J6
kDa
15% SDS-PAGE
Coomassie
IMAC SEC
Sy7J6
kDa
α-His6 (1:1,000)
α-mouse-HRP (1:10,000)

**Figure S4 | Protein purifications.**

(**A**) *C. jejuni* (*Cj*) MlaC, (**B**) delipidated *Cj* MlaC, (**C**) *E. coli* (*Ec*) MlaC and (**D**) the sybody Sy7J6were purified via immobilized metal affinity chromatography (IMAC) and buffer-exchanged by size exclusion chromatography (SEC). The SEC profile was monitored by measuring absorbance at 280 nm ($A_{280}$) across the elution volume ($V_e$). Purification samples, including the lysate (Lys, 5 µL), flow-through (FT, 5 µL), wash fractions 1–2 (W1–2, 10 µL), and IMAC and SEC eluates (5 µg), were analyzed by Coomassie staining. In addition, the His-tagged proteins in the IMAC and SEC eluates were detected by Western blot using the antibodies as indicated. The Western blots shown in (**C**) and (**D**) were cropped as indicated by the black vertical line.

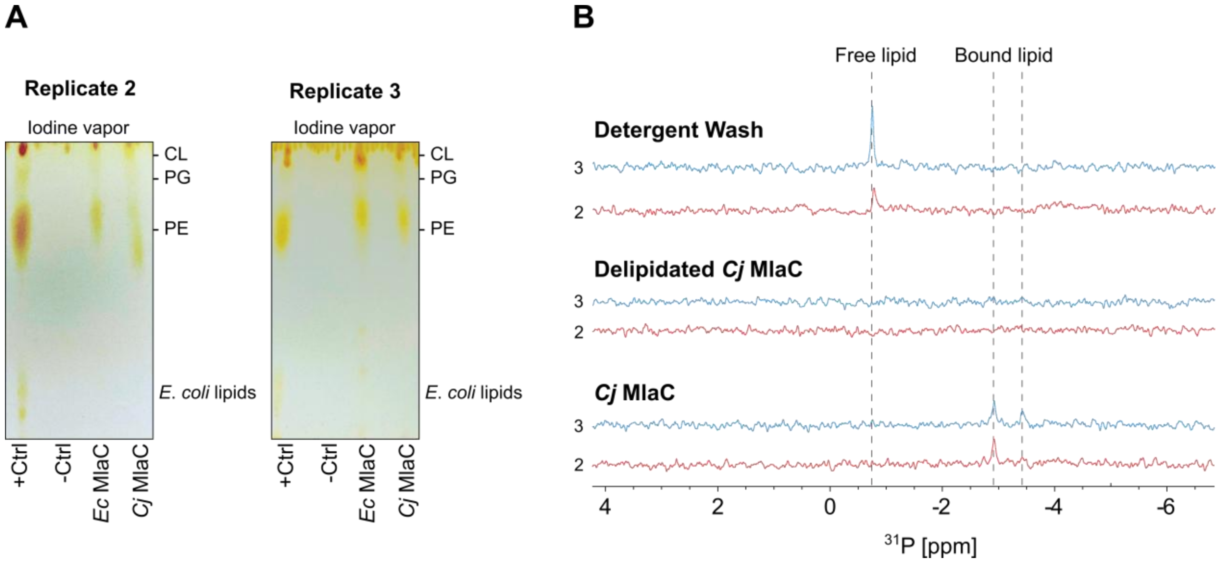


**Figure S5 | Lipid preference of MlaC.**

**(A)** Phospholipids co-purified with *C. jejuni* (*Cj*) and *E. coli* (*Ec*) MlaC were extracted and analyzed by thin-layer chromatography (TLC). Lipids were separated on TLC plates using a chloroform:methanol:acetic acid solvent system (65:25:10, v/v/v) and visualized by staining with iodine vapor. Total lipid extracts from *E. coli* served as a positive control (+Ctrl), while lipids extracted from a periplasmically expressed sybody were used as negative control (-Ctrl). The major *E. coli* phospholipid (PL) species were identified based on migration relative to the positive control and include phosphatidylethanolamine (PE), phosphatidylglycerol (PG) and cardiolipin (CL). (**B**) $^{31}$P NMR analysis on purified *Cj* MlaC, delipidated *Cj* MlaC, and the detergent wash fraction to assess whether the co-purified lipids represent phospholipids. This is the expanded dataset for Figure 2A and B showing the remaining biological duplicates.

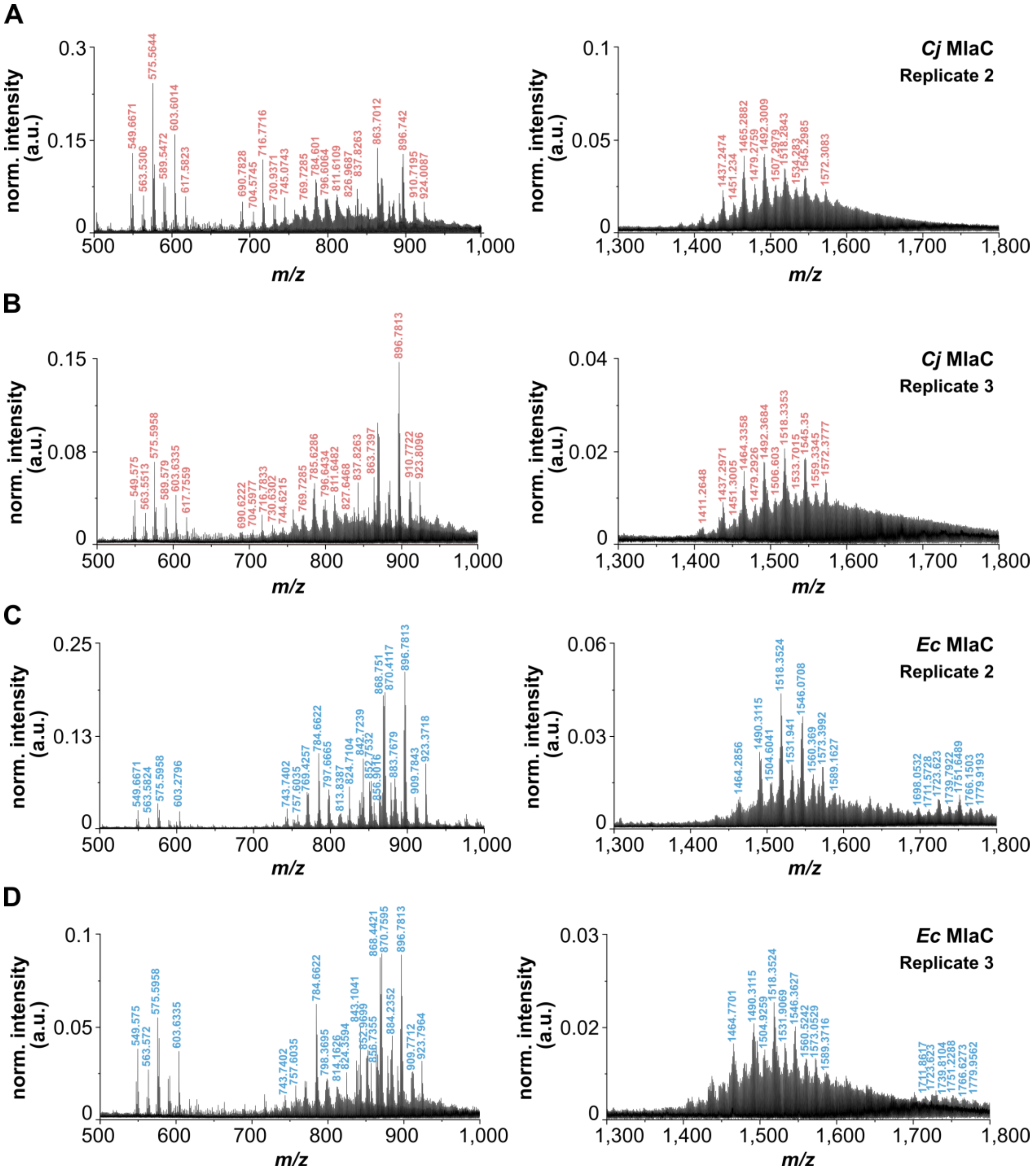


**Figure S6 | nESI-MS analysis of lipid extracts from MlaC.**

(**A-B**) Lipid extracts from purified *Cj* and (**C-D**) *Ec* MlaC were analyzed by nano-electrospray ionization mass spectrometry (nESI-MS) to identify co-purified phospholipids. Peaks that were unambiguously assigned to one of the major *E. coli* phospholipid species using the MAPS® lipid database and that were consistently detected in all three biological replicates were marked and listed in Table S3. This is the expanded dataset for Figure 2D-G showing the remaining biological duplicates.

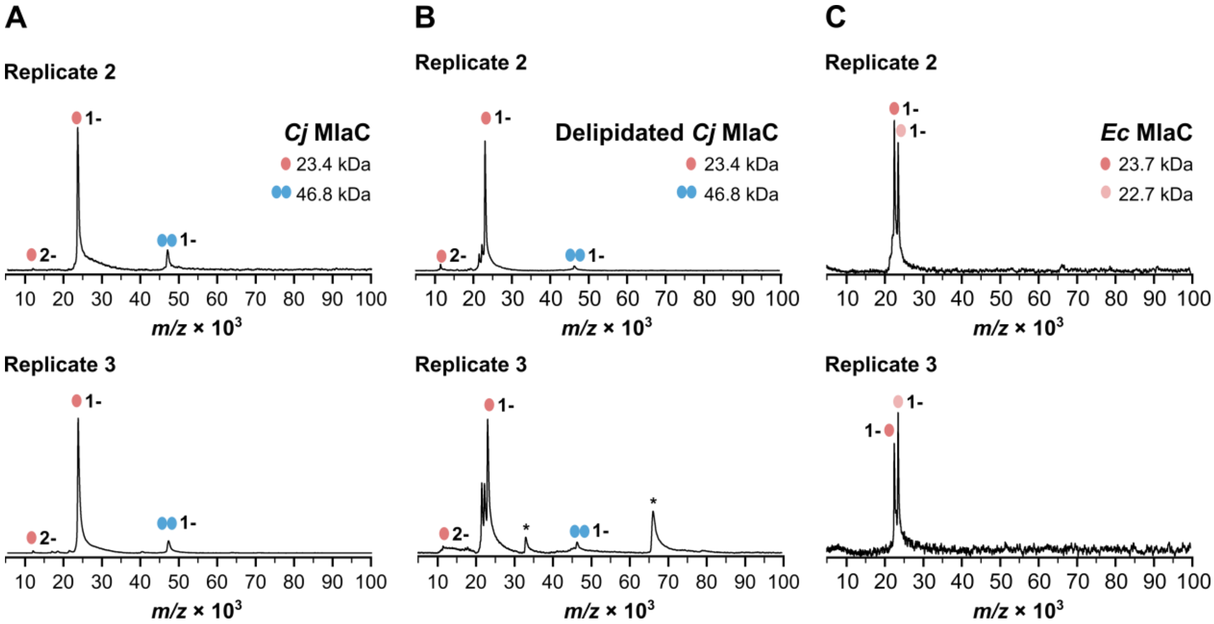


**Figure S7 | LILBID-MS analysis.**

(**A**) Laser-induced liquid bead ion desorption mass spectrometry (LILBID-MS) analysis of purified *C. jejuni* (*Cj*) MlaC, (**B**) delipidated *Cj* MlaC, and (**C**) *E. coli* (*Ec*) MlaC. This is the expanded dataset for Figure 3A showing the remaining biological duplicates. Peaks were assigned to monomers (red) or dimers (blue) in different charge states (**A**, **B**), or to the signal-peptide-containing form and cleaved form of *Ec* MlaC (C) as indicated. *Cj* MlaC was expressed without the signal peptide while *Ec* MlaC was expressed with the signal peptide. The peaks marked with an asterisk in the third replicate of the delipidated *Cj* MlaC sample correspond to an impurity.

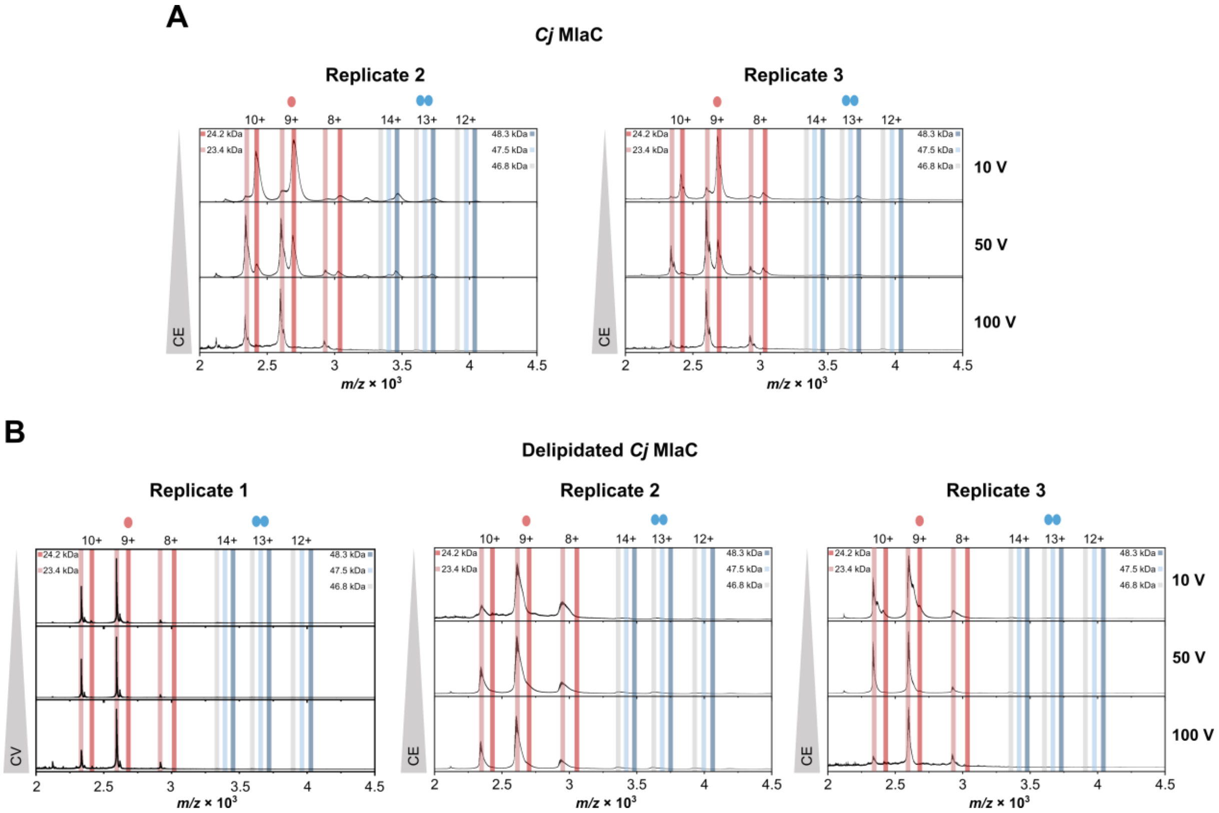


**Figure S8 | nESI-MS analysis of *Cj* MlaC.**

(**A, B**) Nano-electrospray ionization mass spectrometry (nESI-MS) analysis of endogenously lipidated or delipidated *Cj* MlaC at collision energies (CE) of 10 V, 50 V and 100 V. Peaks were assigned to the monomeric (red) or dimeric (blue, gray) conformation in different charge states. This is the expanded dataset for Figure 3B showing the remaining biological duplicates.

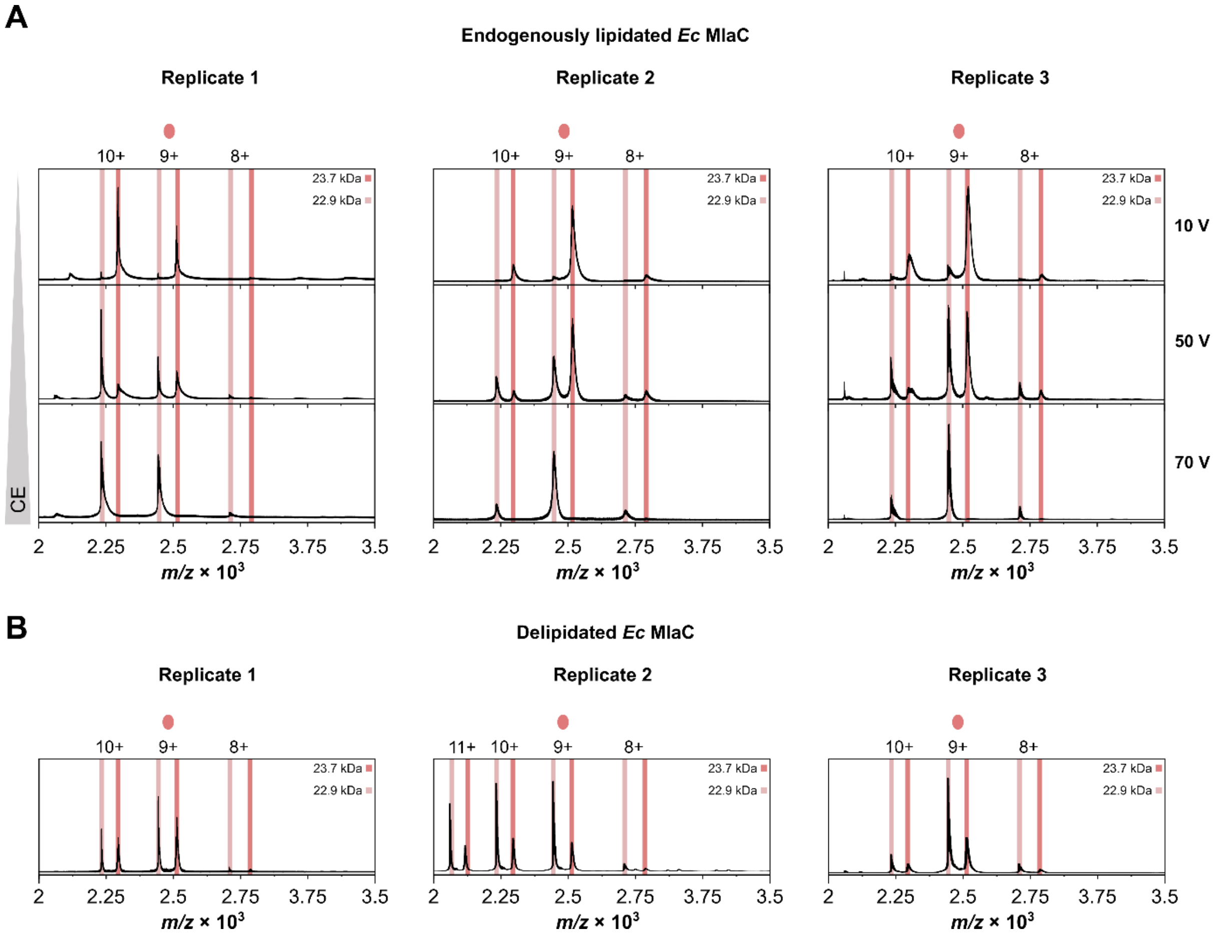


**Figure S9 | nESI-MS analysis of *Ec* MlaC.**

(**A**) Nano-electrospray ionization mass spectrometry (nESI-MS) analysis of purified *Ec* MlaC with endogenously bound lipids at different collision energies (CE). (**B**) nESI-MS spectra of delipidated *Ec* MlaC by *n*-nonyl-β-D-glucopyranoside treatment at 10 V. Peaks were assigned to the lipid-bound (dark red) or lipid-free (light red) state in different charge states

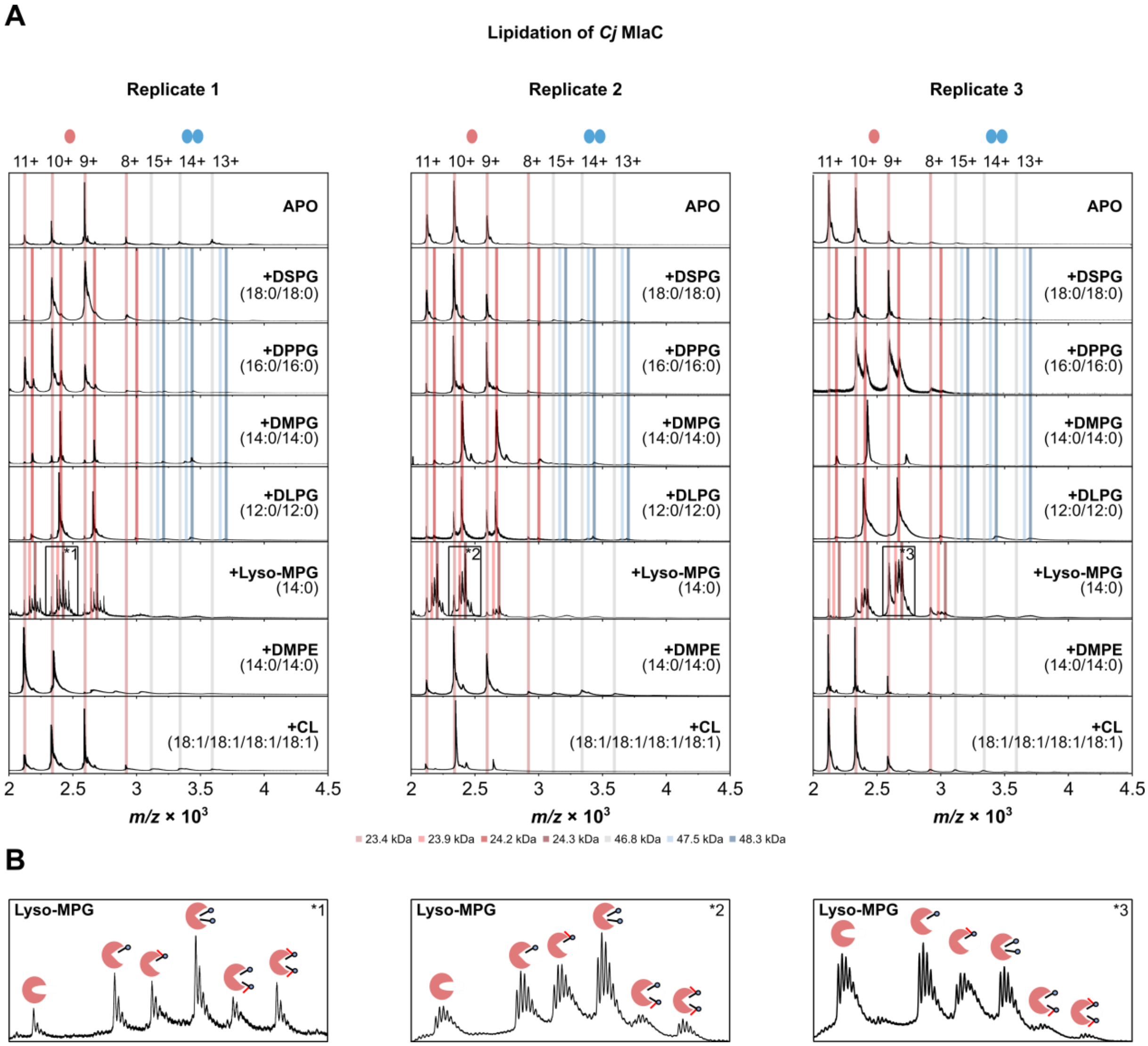


**Figure S10 | Lipidation of *Cj* MlaC *in vitro*.**

(**A**) nESI-MS analysis at 10 V of delipidated *Cj* MlaC in the apo form or after incubation with the phospholipids listed in Table S2. Peaks were assigned to the monomeric (red) or dimeric (blue, gray) conformation in different charge states. (**B**) Zoom-in of the lyso-MPG-bound *Cj* MlaC spectra as indicated by the rectangles in (**A**) and its potential binding behavior. Besides the peaks corresponding to the binding of one or two lyso-MPG molecules, three additional species were detected that could not be unambiguously assigned. These species may result from the binding of a side product generated during lipid synthesis or storage that carries an additional mass of 156.2 Da and is covalently attached to the lyso-MPG molecule.

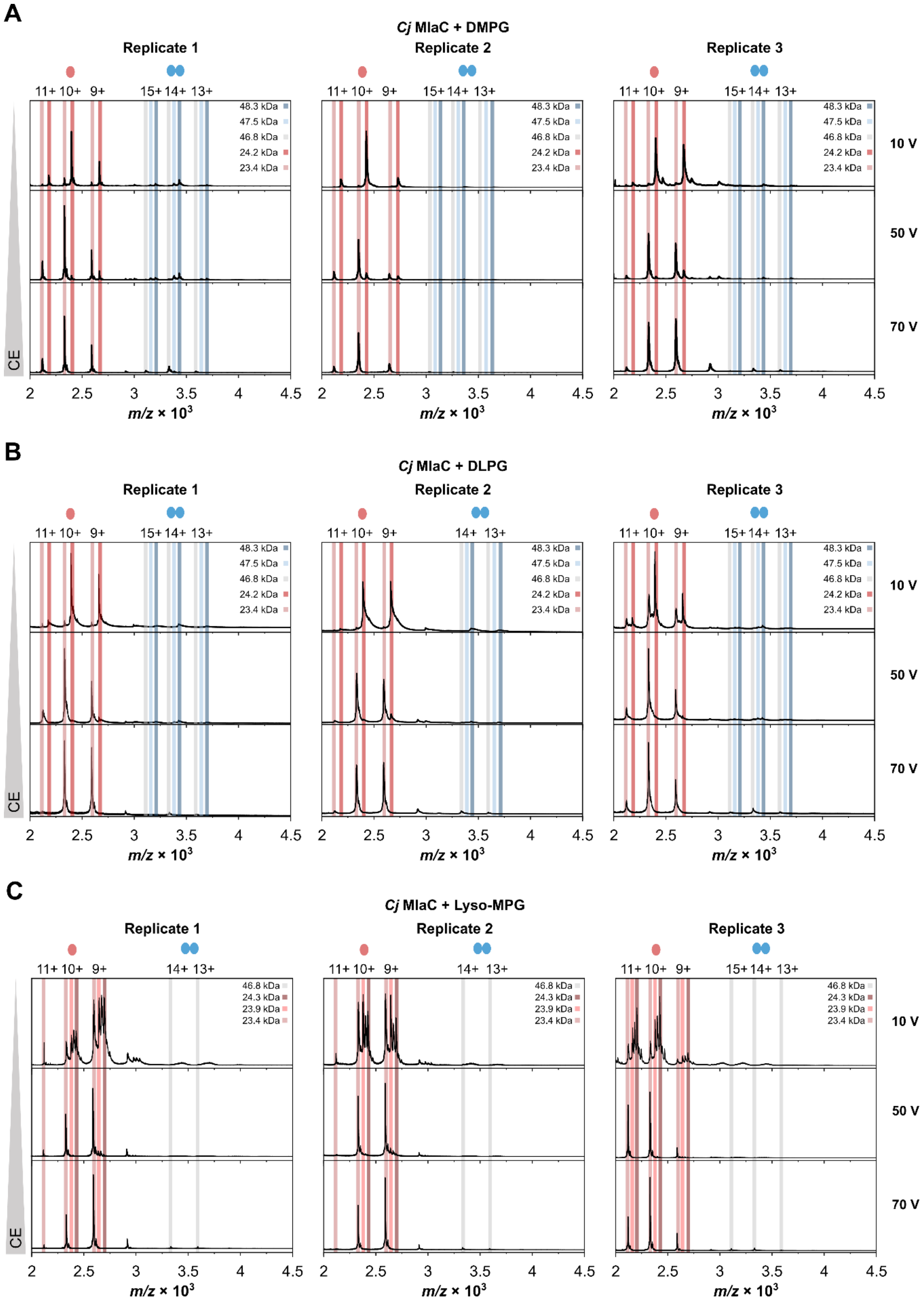


**Figure S11 | nESI-MS analysis of *Cj* MlaC loaded with DMPG and DLPG.**

(**A**) Nano-electrospray ionization mass spectrometry (nESI-MS) analysis of purified *Cj* MlaC with bound 1,2-dimyristoyl-sn-glycero-3-phospho-(1′*rac*glycerol) (DMPG), (**B**) 1,2-didodecanoyl-*sn*-glycero-3-phospho-(1′*rac*glycerol) (DLPG) or (**C**) 1-myristoyl-2-

hydroxy-sn-glycero-3-phospho-(1′*rac*glycerol) (Lyso-MPG) at different collision energies (CE). Purified *Cj* MlaC was delipidated with *n*-nonyl-β-D-glucopyranoside and loaded *in vitro* with DMPG or DLPG. Peaks were assigned to the monomeric (red) or dimeric (blue, gray) conformation in different charge states.

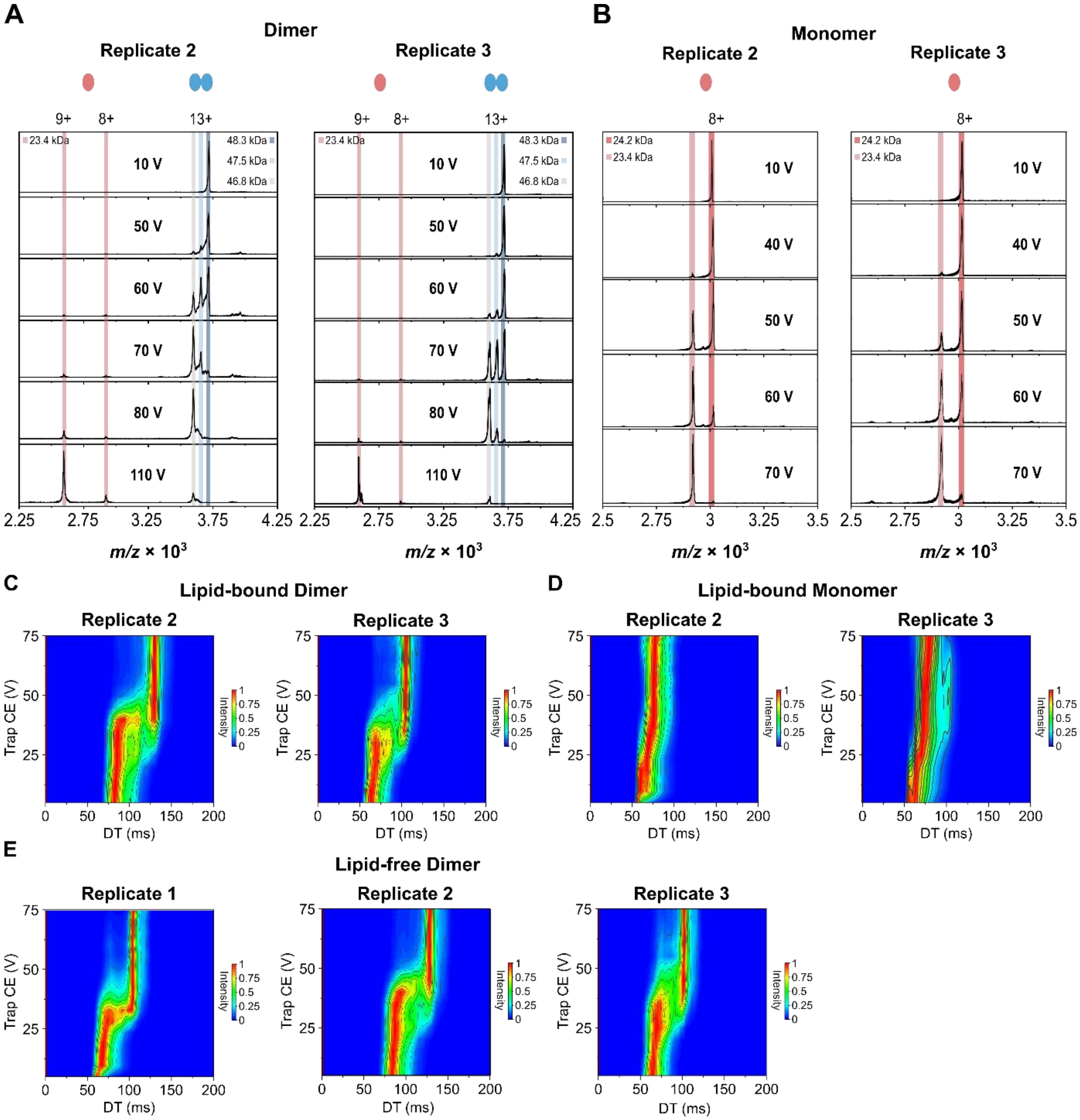


**Figure S12 | Conformational dynamics of *Cj* MlaC during lipid and dimer dissociation**.

(**A**) Tandem mass spectrometry (MS/MS) analysis at different collision voltages filtered for masses corresponding to the *Cj* MlaC dimer (13+) or (**B**) monomer (8+) with bound lipids, respectively. (**C**) Collision-induced unfolding ion mobility spectrometry (CIU IMS) analysis of the lipid-bound dimer peak (13+), (**D**) monomer peak (8+), and (**E**) lipid-free dimer peak (13+), respectively. The trap collision energy (CE) was plotted as a function of the drift time (DT). The spectra shown here are biological replicates of those in Figure 4B.

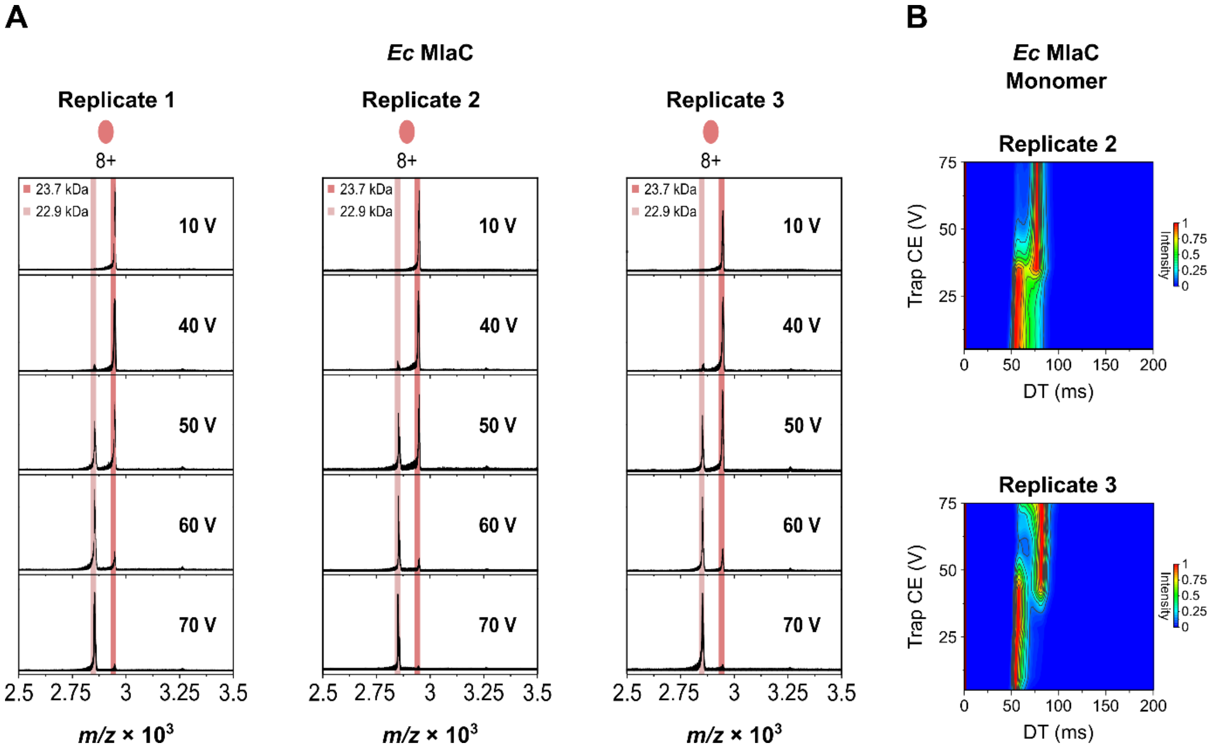


**Figure S13 | Conformational dynamics of *Ec* MlaC during lipid dissociation**.
(**A**) Tandem mass spectrometry (MS/MS) analysis at different collision voltages filtered for masses corresponding to the *Ec* MlaC monomer (8+) with a bound lipid. (**B**) Collision-induced unfolding ion mobility spectrometry (CIU IMS) analysis of the monomer peak (8+). The trap collision energy (CE) was plotted as a function of the drift time (DT). The trace maps shown here are biological replicates of those in Figure 4B.

**Table S1 | Cloning primers used in this study.**

| Primer | Sequence (5'→ 3') |
|---|---|
| *Ec mlaC* fw (NdeI) | TATATACATATGGCAGACCAGACCAATCC |
| *Ec mlaC* rev (XhoI) | ATATCTCGAG TTTTTTCTCTTCCAGAGTGATTTT |
| pET24a_N-term His6_rev | GTGGTGGTGGTGGTGGTGCATATGTATATCTCCTTCTTAAAGTTAAAC |
| *Cj mlaC*+N-term His6 fw | ATGCACCACCACCACCACCACCTCGAGTTACAACTAGATGAAATTTCTAGC |
| *Cj mlaC* with Avi fw | CGAGGCCCAGAAGATCGAGTGGCACGAGTGAGATCCGGCTGCTAA |
| *Cj mlaC* with Avi rev | AAGATATCGTTCAGGCCGCCGCTGCT TTCTATGACAATACTTTCAAGTTTT |
| pTTQ18_*sfGFP*_fw | CGGATGAGCTCTACAAAACTAGTCATCACCATCATCATCATCA |
| *Cj mlaC_sfGFP*_rev | GACCTTGAAACAAAACTTCTAATTCTATGACAATACTTTCAAGTTTTT |
| *sfGFP* fw | TTAGAAGTTTTGTTTCAAGGTC |

| | |
|---|---|
| *sfGFP* rev | ACTAGTTTTGTAGAGCTCATCCG |

**Table S2 | Phospholipids used in this study.**

| Lipid name | Abbreviation | Supplier |
|---|---|---|
| 1,1′,2,2′-tetra-(9Z-octadecenoyl) cardiolipin | CL | Avanti Polar Lipids, 710335P |
| 1,2-didodecanoyl-*sn*-glycero-3-phospho-(1′*rac*glycerol) | DLPG | Avanti Polar Lipids, 840435P |
| 1,2-ditetradecyl-rac-glycero-3-phosphoethanolamine | DMPE | MedChemExpress, HY-142983 |
| 1,2-dimyristoyl-sn-glycero-3-phospho-(1′*rac*glycerol) | DMPG | Avanti Polar Lipids, 840445P |
| 1,2-di-(9Z-octadecenoyl)-*sn*-glycero-3-phosphoethanolamine | DOPE | Avanti Polar Lipids, 850725P |
| 1,2-di-(9Z-octadecenoyl)-*sn*-glycero-3-phospho-(1′*rac*glycerol) | DOPG | Avanti Polar Lipids, 840475P |
| 1,2-dipalmitoyl-sn-glycero-3-phosphoglycerol | DPPG | MedChemExpress, HY-125940 |
| 1,2-dioctadecanoyl-*sn*-glycero-3-phospho-(1′*rac*glycerol) | DSPG | Avanti Polar Lipids, 840465P |
| 1-myristoyl-2-hydroxy-sn-glycero-3-phospho-(1′*rac*glycerol) | Lyso-MPG | Avanti Polar Lipids, 858120P |

**Table S3 | Co-purified phospholipids from *C. jejuni* and *E. coli* MlaC.**

The most abundant phospholipid species that consistently co-purified with *C. jejuni* and *E. coli* MlaC across all biological replicates. Only phospholipid species that were reproducibly detected and could be unambiguously assigned to one of the major *E. coli* phospholipid classes were included. Lipid identification was performed using the Lipid MAPS® database [33,34].

| ***C. jejuni* MlaC** | | | ***E. coli* MlaC** | | |
|---|---|---|---|---|---|
| **Input Mass** | **Name** | **Formula** | **Input Mass** | **Name** | **Formula** |
| 549.67 | Lyso PG 19:0 | $[M+Na]^+$ | 549.36 | Lyso PG 19:0 | $[M+Na]^+$ |
| 563.53 | Lyso PG 20:0 | $[M+Na]^+$ | 743.75 | PG 34:4 | $[M+H]^+$ |
| 589.55 | Lyso PG 22:1 | $[M+Na]^+$ | | PG 32:1 | $[M+Na]^+$ |
| 690.78 | PE 32:1 | $[M+H]^+$ | | PE 35:4 | $[M+NH_4]^+$ |

| | | | | | |
|---|---|---|---|---|---|
| 704.57 | PE 33:1 | $[M+H]^+$ | 757.62 | PG 35:4 | $[M+H]^+$ |
| | PE 32:5 | $[M+Na]^+$ | | PG 33:1 | $[M+Na]^+$ |
| | PG 30:4 | $[M+NH_4]^+$ | | PE 36:4 | $[M+NH_4]^+$ |
| 716.77 | PE 34:2 | $[M+H]^+$ | 769.43 | PG 36:5 | $[M+H]^+$ |
| 730.94 | PE 35:2 | $[M+H]^+$ | | PG 34:2 | $[M+Na]^+$ |
| | PE 34:6 | $[M+Na]^+$ | | PE 37:5 | $[M+NH_4]^+$ |
| | PG 32:5 | $[M+NH_4]^+$ | | CL 74:0 | $[M+2Na]^{2+}$ |
| 745.07 | PG 34:3 | $[M+H]^+$ | 784.66 | PE 39:3 | $[M+H]^+$ |
| | PG 32:0 | $[M+Na]^+$ | | PE 40:10 | $[M+H]^+$ |
| | PE 35:3 | $[M+NH_4]^+$ | | PE 37:0 | $[M+Na]^+$ |
| 769.73 | PG 36:5 | $[M+H]^+$ | | PE 38:7 | $[M+Na]^+$ |
| | PG 34:2 | $[M+Na]^+$ | | PG 36:6 | $[M+NH_4]^+$ |
| | PE 37:5 | $[M+NH_4]^+$ | | CL 78:13 | $[M+2Na]^{2+}$ |
| | CL 74:0 | $[M+2Na]^{2+}$ | 797.67 | PG 38:5 | $[M+H]^+$ |
| 784.60 | PE 39:3 | $[M+H]^+$ | | PG 36:2 | $[M+Na]^+$ |
| | PE 40:10 | $[M+H]^+$ | | PE 39:5 | $[M+NH_4]^+$ |
| | PE 37:0 | $[M+Na]^+$ | | CL 78:0 | $[M+2Na]^{2+}$ |
| | PE 38:7 | $[M+Na]^+$ | 814.03 | PG 39:4 | $[M+H]^+$ |
| | PG 36:6 | $[M+NH_4]^+$ | | PG 37:1 | $[M+Na]^+$ |
| | CL 78:13 | $[M+2Na]^{2+}$ | | PE 40:4 | $[M+NH_4]^+$ |
| 796.61 | PE 40:4 | $[M+H]^+$ | 824.33 | PE 42:4 | $[M+H]^+$ |
| | PE 38:1 | $[M+Na]^+$ | | PE 40:1 | $[M+Na]^+$ |
| | PE 39:8 | $[M+Na]^+$ | | PG 39:7 | $[M+NH_4]^+$ |
| | PG 36:0 | $[M+H]^+$ | | PG 38:0 | $[M+NH_4]^+$ |
| | PG 37:7 | $[M+NH_4]^+$ | 842.72 | PE 43:2 | $[M+H]^+$ |
| | CL 78:1 | $[M+2Na]^{2+}$ | | PE 42:6 | $[M+Na]^+$ |
| 811.61 | PG 39:5 | $[M+H]^+$ | | PG 40:5 | $[M+NH_4]^+$ |

| | | | | | |
|---|---|---|---|---|---|
| | PG 37:2 | $[M+Na]^+$ | 852.75 | PE 44:4 | $[M+H]^+$ |
| | PG 38:9 | $[M+Na]^+$ | | PE 42:1 | $[M+Na]^+$ |
| | PE 40:5 | $[M+NH_4]^+$ | | PG 40:0 | $[M+NH_4]^+$ |
| | CL 80:0 | $[M+2Na]^{2+}$ | | PG 41:7 | $[M+NH_4]^+$ |
| 826.97 | PE 42:3 | $[M+H]^+$ | 856.59 | PE 44:2 | $[M+H]^+$ |
| | PE 40:0 | $[M+Na]^+$ | | PE 43:6 | $[M+Na]^+$ |
| | PE 41:7 | $[M+Na]^+$ | | PG 41:5 | $[M+NH_4]^+$ |
| | PG 39:6 | $[M+NH_4]^+$ | 868.75 | PE 43:0 | $[M+Na]^+$ |
| 837.83 | PG 41:6 | $[M+H]^+$ | | PE 44:7 | $[M+Na]^+$ |
| | PG 39:3 | $[M+Na]^+$ | | PG 42:6 | $[M+NH_4]^+$ |
| | PG 40:10 | $[M+Na]^+$ | 870.76 | PE 44:6 | $[M+Na]^+$ |
| | PE 42:6 | $[M+NH_4]^+$ | | PG 42:5 | $[M+NH_4]^+$ |
| 863.70 | PG 42:0 | $[M+H]^+$ | 883.87 | PG 44:4 | $[M+H]^+$ |
| | PG 41:4 | $[M+Na]^+$ | | PG 42:1 | $[M+Na]^+$ |
| | PG 42:11 | $[M+Na]^+$ | | PG 44:6 | $[M+NH_4]^+$ |
| | PE 43:0 | $[M+NH_4]^+$ | 909.56 | PG 44:2 | $[M+Na]^+$ |
| | PE 44:7 | $[M+NH_4]^+$ | 1464.14 | CL 72:1 | $[M+H]^+$ |
| 896.74 | PG 44:6 | $[M+NH_4]^+$ | 1490.31 | CL 74:2 | $[M+H]^+$ |
| 1451.23 | CL 70:2 | $[M+NH_4]^+$ | 1504.62 | CL 74:3 | $[M+NH_4]^+$ |
| 1479.28 | CL 72:2 | $[M+NH_4]^+$ | 1518.35 | CL 76:2 | $[M+H]^+$ |
| 1492.30 | CL 74:1 | $[M+H]^+$ | 1531.94 | CL 78:9 | $[M+H]^+$ |
| 1507.30 | CL 74:2 | $[M+NH_4]^+$ | | CL 76:6 | $[M+Na]^+$ |
| 1518.28 | CL 76:2 | $[M+H]^+$ | 1546.07 | CL 80:16 | $[M+H]^+$ |
| 1534.28 | CL 78:8 | $[M+H]^+$ | | CL 78:2 | $[M+H]^+$ |
| 1534.28 | CL 76:5 | $[M+Na]^+$ | | CL 78:13 | $[M+Na]^+$ |
| 1572.31 | CL 78:0 | $[M+Na]^+$ | | | |

**Table S4| Calculated lipidation ratios of MlaC.**

Ratios of lipid-bound MlaC compared to lipid free MlaC. Ratios were determined by the summing of all peak integrals in one spectrum corresponding to lipid-bound MlaC, divided by the sum of the peak integrals of lipid free MlaC.

| Protein | Lipid | Repeat | CE | Ratio lipid bound/ unbound |
|---|---|---|---|---|
| *Cj* MlaC Monomer | Endogenous | 1 | 10 | 0.74 |
| | | 2 | 10 | 0.83 |
| | | 3 | 10 | 0.86 |
| | | **Average** | **10** | **0.81 ± 0.05** |
| *Cj* MlaC Dimer | Endogenous | 1 | 10 | 0.86 |
| | | 2 | 10 | 0.97 |
| | | 3 | 10 | 0.92 |
| | | **Average** | **10** | **0.91 ± 0.05** |
| *Cj* MlaC Monomer | Endogenous | 1 | 50 | 0.44 |
| | | 2 | 50 | 0.38 |
| | | 3 | 50 | 0.38 |
| | | **Average** | **50** | **0.40 ± 0.03** |
| *Cj* MlaC Dimer | Endogenous | 1 | 50 | 0.83 |
| | | 2 | 50 | 0.81 |
| | | 3 | 50 | 0.92 |
| | | **Average** | **50** | **0.85 ± 0.05** |
| *Ec* MlaC Monomer | Endogenous | 1 | 10 | 0.96 |
| | | 2 | 10 | 0.91 |
| | | 3 | 10 | 0.86 |
| | | **Average** | **10** | **0.90 ± 0.05** |
| *Ec* MlaC Monomer | Endogenous | 1 | 50 | 0.52 |
| | | 2 | 50 | 0.58 |
| | | 3 | 50 | 0.52 |
| | | **Average** | **50** | **0.54 ± 0.04** |

| | | | | |
|---|---|---|---|---|
| *Cj* MlaC Monomer | DMPG *in vitro* | 1 | 10 | 0.85 |
| | | 2 | 10 | 0.94 |
| | | 3 | 10 | 0.92 |
| | | **Average** | **10** | **0.90 ± 0.04** |
| *Cj* MlaC Dimer | DMPG *in vitro* | 1 | 10 | 0.95 |
| | | 2 | 10 | 0.81 |
| | | 3 | 10 | 0.84 |
| | | **Average** | **10** | **0.87 ± 0.06** |
| *Cj* MlaC Monomer | DMPG *in vitro* | 1 | 50 | 0.12 |
| | | 2 | 50 | 0.27 |
| | | 3 | 50 | 0.23 |
| | | **Average** | **50** | **0.21 ± 0.07** |
| *Cj* MlaC Dimer | DMPG *in vitro* | 1 | 50 | 0.91 |
| | | 2 | 50 | 0.76 |
| | | 3 | 50 | 0.93 |
| | | **Average** | **50** | **0.87 ± 0.07** |
| *Cj* MlaC Monomer | DLPG *in vitro* | 1 | 10 | 0.94 |
| | | 2 | 10 | 0.66 |
| | | 3 | 10 | 0.92 |
| | | **Average** | **10** | **0.84 ± 0.13** |
| *Cj* MlaC Dimer | DLPG *in vitro* | 1 | 10 | 0.88 |
| | | 2 | 10 | 0.84 |
| | | 3 | 10 | 0.90 |
| | | **Average** | **10** | **0.87 ± 0.03** |
| *Cj* MlaC Monomer | DLPG *in vitro* | 1 | 50 | 0.10 |
| | | 2 | 50 | 0.12 |
| | | 3 | 50 | 0.20 |

| | | | | |
|---|---|---|---|---|
| | | **Average** | **50** | **0.14 ± 0.05** |
| *Cj* MlaC Dimer | DLPG *in vitro* | 1 | 50 | 0.86 |
| | | 2 | 50 | 0.77 |
| | | 3 | 50 | 0.83 |
| | | **Average** | **50** | **0.82 ± 0.04** |
| *Cj* MlaC Monomer | Lyso-MPG *in vitro* | 1 | 10 | 0.75 |
| | | 2 | 10 | 0.92 |
| | | 3 | 10 | 0.64 |
| | | **Average** | **10** | **0.77 ± 0.12** |
| *Cj* MlaC Dimer | Lyso-MPG *in vitro* | 1 | 10 | 0.90 |
| | | 2 | 10 | 0.83 |
| | | 3 | 10 | 0.85 |
| | | **Average** | **10** | **0.86 ± 0.10** |
| *Cj* MlaC Monomer | Lyso-MPG *in vitro* | 1 | 50 | 0.19 |
| | | 2 | 50 | 0.25 |
| | | 3 | 50 | 0.27 |
| | | **Average** | **50** | **0.23 ± 0.04** |
| *Cj* MlaC Dimer | Lyso-MPG *in vitro* | 1 | 50 | 0.86 |
| | | 2 | 50 | 0.85 |
| | | 3 | 50 | 0.85 |
| | | **Average** | **50** | **0.85 ± 0.01** |
| *Ec* MlaC Monomer | Detergent-delipidated | 1 | 10 | 0.58 |
| | | 2 | 10 | 0.4 |
| | | 3 | 10 | 0.43 |
| | | **Average** | **10** | **0.47 ± 0.10** |
| *Ec* MlaC Monomer | Detergent-delipidated | 1 | 50 | 0.13 |
| | | 2 | 50 | 0.10 |

| | | | | |
|---|---|---|---|---|
| | | 3 | 50 | 0.09 |
| | | **Average** | **50** | **0.10 ± 0.02** |

**Table S5| Calculated delipidation efficiencies of MlaC.**

Ratios of lipid bound MlaC compared to lipid free MlaC. Ratios were determined by the summing of all peak integrals in one spectrum corresponding to lipid-bound MlaC, divided by the sum of the peak integrals of lipid free MlaC.

| Lipid extract | Repeat | CE | Ratio Lyso-PGs/all PGs |
|---|---|---|---|
| *Cj* MlaC | 1 | 10 | 7.4% |
| | 2 | 10 | 6.6% |
| | 3 | 10 | 8.0% |
| | **Average** | **10** | **7.4 ± 0.6%** |
| *Ec* MlaC | 1 | 10 | 2.5% |
| | 2 | 10 | 1.6% |
| | 3 | 10 | 3.4% |
| | **Average** | **10** | **2.4 ± 0.8%** |
